# In mice, discrete odors can selectively promote the neurogenesis of sensory neuron subtypes that they stimulate

**DOI:** 10.1101/2024.02.10.579748

**Authors:** Kawsar Hossain, Madeline Smith, Karlin E. Rufenacht, Rebecca O’Rourke, Stephen W. Santoro

## Abstract

In mammals, olfactory sensory neurons (OSNs) are born throughout life, ostensibly solely to replace neurons lost *via* turnover or injury. This assumption follows from the hypothesis that olfactory neurogenesis is stochastic with respect to neuron subtype, as defined by the single odorant receptor that each neural precursor stochastically chooses out of hundreds of possibilities. This assumption is challenged, however, by recent findings that the birthrates of a fraction of OSN subtypes are selectively reduced by olfactory deprivation. These findings raise questions about how, and why, olfactory stimuli are required to accelerate the neurogenesis rates of some subtypes, including whether the stimuli are specific (e.g., discrete odorants) or generic (e.g., broadly activating odors or mechanical stimuli). Based on previous findings that the exposure of mice to sex-specific odors can increase the representations of subtypes responsive to those odors, we hypothesized that the neurogenic stimuli comprise discrete odorants that selectively stimulate OSNs of the same subtypes whose birthrates are accelerated. In support of this, we have found, using scRNA-seq and subtype-specific OSN birthdating, that exposure to male and exogenous musk odors can accelerate the birthrates of subtypes responsive to those odors. These findings reveal that certain odor experiences can selectively ‘amplify’ specific OSN subtypes and suggest that persistent OSN neurogenesis serves, in part, an adaptive function.

## Introduction

Mammalian olfactory epithelia (OE) contain hundreds of distinct olfactory sensory neuron (OSN) subtypes, each of which expresses a single odorant receptor (OR) and can thereby be stimulated and/or inhibited by a distinct set of odorant molecules ^1,2^. The olfactory epithelium is one of a few regions of the mammalian nervous system where neurogenesis occurs throughout life ^3–5^. In the hippocampus and olfactory bulb, persistent neurogenesis plays vital roles in learning and memory ^6–8^. By contrast, life-long neurogenesis within the mammalian OE is generally assumed to function solely to replace OSNs that are lost due to normal turnover or environmentally induced damage. This assumption is consistent with the prevailing hypothesis that OSN neurogenesis is strictly stochastic with respect to subtype since it is based on the evidently stochastic process of OR choice ^9–11^.

Several studies have observed that the relative quantities of distinct OSN subtypes can be altered by manipulating olfactory experience in mice ^12–24^. Olfactory deprivation on one side of the OE *via* unilateral naris occlusion (UNO), for example, has been found to cause both increases and decreases in the representations of different OSN subtypes on the closed side relative to the open ^21,23^. Similar bidirectional changes in subtype representations have been observed following olfactory enrichment *via* exposure to discrete odors in mice ^17,19,24^. Experience-induced changes in the representations of specific OSN subtypes have generally been attributed entirely to altered OSN survival ^12,13,18–26^, which could theoretically account for both increases and decreases in subtype representations. By contrast, the stochastic nature of OR choice would seem to preclude the contribution of a mechanism involving subtype-selective changes in OSN birthrates.

Unexplained observations that olfactory deprivation *via* UNO reduces both the overall rate of OSN neurogenesis ^27–30^ and the representations of a fraction of OSN subtypes ^14,16,19,21,23,30^ motivated a recent study to test the possibility that the birthrates of these subtypes depend on olfactory stimulation. To do so, newborn OSNs of subtypes that exhibit altered representations following UNO were quantified on the open side relative to the closed side of the OEs from UNO-treated mice ^30^. Remarkably, subtypes previously found to have reduced representations on the closed side of the OE also exhibited significantly lower birthrates on the closed side, while subtypes with elevated representations on the closed side showed no differences in birthrates between the two sides. These findings indicated that olfactory stimulation affects the representations of specific OSN subtypes *via* two distinct mechanisms. One mechanism reduces the representations of specific subtypes on the closed side of the OE *via* selective reductions in their birthrates as a result of olfactory deprivation. By contrast, a second mechanism elevates on the closed side the representations of subtypes with extremely high levels of baseline activity, presumably *via* lengthened OSN lifespans due to protection from overstimulation ^30^.

Findings that the birthrates of a fraction of subtypes depend on olfactory stimulation raise several fascinating questions related to the mechanism and function of this phenomenon. One key question concerns the nature of the stimuli that promote neurogenesis. Because naris occlusion reduces exposure to potentially thousands of odors, as well as mechanical stimuli, and likely causes additional physiological changes ^31^, we envisioned that the neurogenic stimuli could be either non-specific with respect to the subtypes whose birthrates are reduced by olfactory deprivation (e.g., generic odors, mechanical stimuli, or other physiological effects of UNO) or, alternatively, discrete odorants that selectively stimulate the same OSN subtypes whose birthrates are affected. If the neurogenic stimuli are non-specific, this would imply a generic mechanism and, perhaps, an unknown homeostatic function. By contrast, if the stimuli are discrete odorants that selectively activate the same subtypes whose birthrates are affected, this would imply a highly specific mechanism in which exposure to certain odors can “amplify” subtypes responsive to those odors.

In this study, we tested the possibility that the neurogenic stimuli comprise discrete odorants. One prediction of this hypothesis is that the extent to which naris occlusion reduces the birthrates of specific subtypes should vary depending on the odor environment to which animals are exposed. Consistent with this, open-side biases in the birthrates of specific subtypes have been found to depend on whether a mouse was in the nursing or post-weaning stage at the time of birthrate assessment ^30^. A second prediction is that exposure of intact (non-occluded) mice to specific odors should selectively increase the representations of subtypes responsive to those odors. Support for this comes from two separate studies that identified a set of subtypes that were differentially represented within the OEs of male and female mice that were housed in a sex-separated manner until six months ^19^ or up to 43 weeks of age ^24^, and that these differences were attenuated in mice that were housed sex-combined. Notably, several of these subtypes were found to selectively respond to sex-specific odors ^19,24^, indicating that the observed differences were at least partly odor-dependent. As with UNO, the differentially represented subtypes fell into two categories: those for which stimulation increased their representations and those for which stimulation had the opposite effect. OSNs in the former category were exemplified by subtype Olfr235, which exhibited a higher representation in mice exposed to males ^19,24^ and selective responsivity to male-emitted odors ^19^, and was previously shown to respond to musk odors ^32–34^, whose emission by mice has not been documented. In analogy to findings from UNO-based studies ^30^, we speculated that these subtypes undergo sex-specific odor-dependent selective birthrate acceleration. Here we present evidence that the neurogenesis rates of Olfr235 and other musk-responsive OSN subtypes are increased in mice exposed to male-specific and musk odors. These findings support the hypothesis that discrete odors can selectively accelerate the birthrates of OSN subtypes that they stimulate and suggest that the function of persistent OSN neurogenesis is not limited to the replacement of neurons lost through damage or normal turnover, but may also enable adaptive changes to the subtype composition of the OSN population.

## Results

### Long-term exposure of mice to male odors is associated with increased representations of musk-responsive OSN subtypes

To test the possibility that discrete odorants can selectively accelerate the birthrates of the OSN subtypes that they stimulate, we sought to identify subtypes that exhibit odor exposure-dependent increases in representations and for which stimulating odors have been identified. A challenge of this approach is that odorant ligands remain unidentified for most OSN subtypes ^35^. To overcome this, we speculated that candidate subtypes might be identified among those previously found to be more highly represented in mice exposed to odors emitted specifically by male or female mice ^19,24^. OSN subtypes in this category, which include Olfr235, were observed to be more highly represented in mice exposed to male conspecifics (sex-separated males; sex-combined males and females) compared to mice isolated from males (sex-separated females) ^19,24^ (Appendix 1–figure 1). These findings, combined with observations that Olfr235 OSNs show selective responsivity to male odors ^19^, suggest that the exposure of mice to one or more odor components specific to males causes an increase in the representation of subtype Olfr235 within the OSN population. Interestingly, Olfr235, which expresses the OR-encoding gene *Or5an11*, belongs to a group of subtypes that express homologous ORs and respond to different musk odorants with varying levels of sensitivity and selectivity ^32,33^. Notably, like Olfr235, other subtypes within this group, which includes Olfr1440 *(Or5an6)*, Olfr1437 *(Or5an1b)*, Olfr1431 *(Or5an9)*, and Olfr1434 *(Or5an1)*, exhibited higher transcript levels in the OEs of mice exposed to male odors compared to their unexposed counterparts (except subtype Olfr1434, whose transcript levels were too low to be accurately assessed) ^19^ (Appendix 1–figure 1). Accordingly, RNA-fluorescent *in situ* hybridization (RNA-FISH) analyses of a subset of these ORs, Olfr235 and Olfr1437, confirmed that the elevated transcript levels observed in mice exposed to male odors reflected greater representations of these subtypes within the OSN population ^19^. By contrast, subtypes Olfr912 *(Or8b48)* and Olfr1295 (*Or4k45)*, which detect the male-specific non-musk odorants 2-(sec-Butyl)-4,5-dihydrothiazole (SBT) and (methylthio)methanethiol (MTMT), respectively ^24^, exhibited lower representations and/or transcript levels in mice exposed to male odors ^19,24^ (Appendix 1–figure 1), possibly reflecting reduced survival due to overstimulation. Taken together, these findings indicate that OSN subtypes that are responsive to musk odors are selectively increased in their representations upon long-term exposure to male mice, consistent with the hypothesis that components of male odors selectively accelerate the birthrates of these subtypes.

### Olfactory deprivation via UNO reduces quantities of newborn OSNs of musk-responsive subtypes in male mice

If the birthrates of musk-responsive OSN subtypes are accelerated by exposure to male odors, quantities of newborn OSNs of these subtypes would be expected to be reduced following olfactory deprivation in male mice. To test this, we analyzed two single-cell RNA sequencing (scRNA-seq) datasets (one generated previously [OE 1] ^30^ and the other as part of this study [OE 2]) corresponding to the open and closed sides of the OEs from male mice that had been UNO- treated at postnatal day (PD) 14, weaned sex-separated at PD 21, and dissected at PD 28 (Figure 1A, B; Figure 1–figure supplement 1A). Within each dataset, newborn OSNs of specific subtypes can be identified based on the co-expression of *Gap43*+ (an established marker of immature OSNs ^36–38)^ and specific OR genes ^30^ (Figure 1C, D; Figure 1–figure supplement 1B, C). Using this approach, we mapped newborn OSNs of each of the five known or putative musk-responsive subtypes within the feature plots corresponding to the open and closed sides of each OE (Figure 1D; Figure 1–figure supplement 1C) and quantified them as a proportion to total OSNs (Figure 1E-*left*; Figure 1–figure supplement 1D) and total cells (Figure 1–figure supplement 1E) within the open and closed data subsets. Remarkably, musk-responsive OSN subtypes exhibited, on average, 3.1-fold greater quantities of newborn OSNs within the open-sides of the OEs compared to the closed, with all subtypes except Olfr1437 showing open-side biases (Figure 1D, E-*left*; Figure 1–figure supplement 1C-E). By contrast, newborn OSNs of 15 randomly chosen subtypes located within approximately the same region of the OE where musk-responsive subtypes reside (canonical zones 2 and 3) ^39^ comprise, on average, nearly equal quantities within the open and closed sides (1.07-fold difference) (Figure 1E-*right*). To verify that the differences in newborn OSN quantities are not due to aberrant OR expression regulation (e.g, co-expression of multiple ORs), we analyzed cell-specific OR expression at three stages of OSN differentiation: immediate neuronal precursor 3 (INP3), immature OSN (iOSN), and mature OSN (mOSN) (Figure 1–figure supplement 2). As expected, individual iOSNs and mOSNs of specific subtypes exhibit robust and singular OR expression on both the open and closed sides of OEs from UNO-treated mice, while INP3 cells co-express low levels of multiple OR transcripts, as observed previously ^38,40–45^.

**Figure 1.**
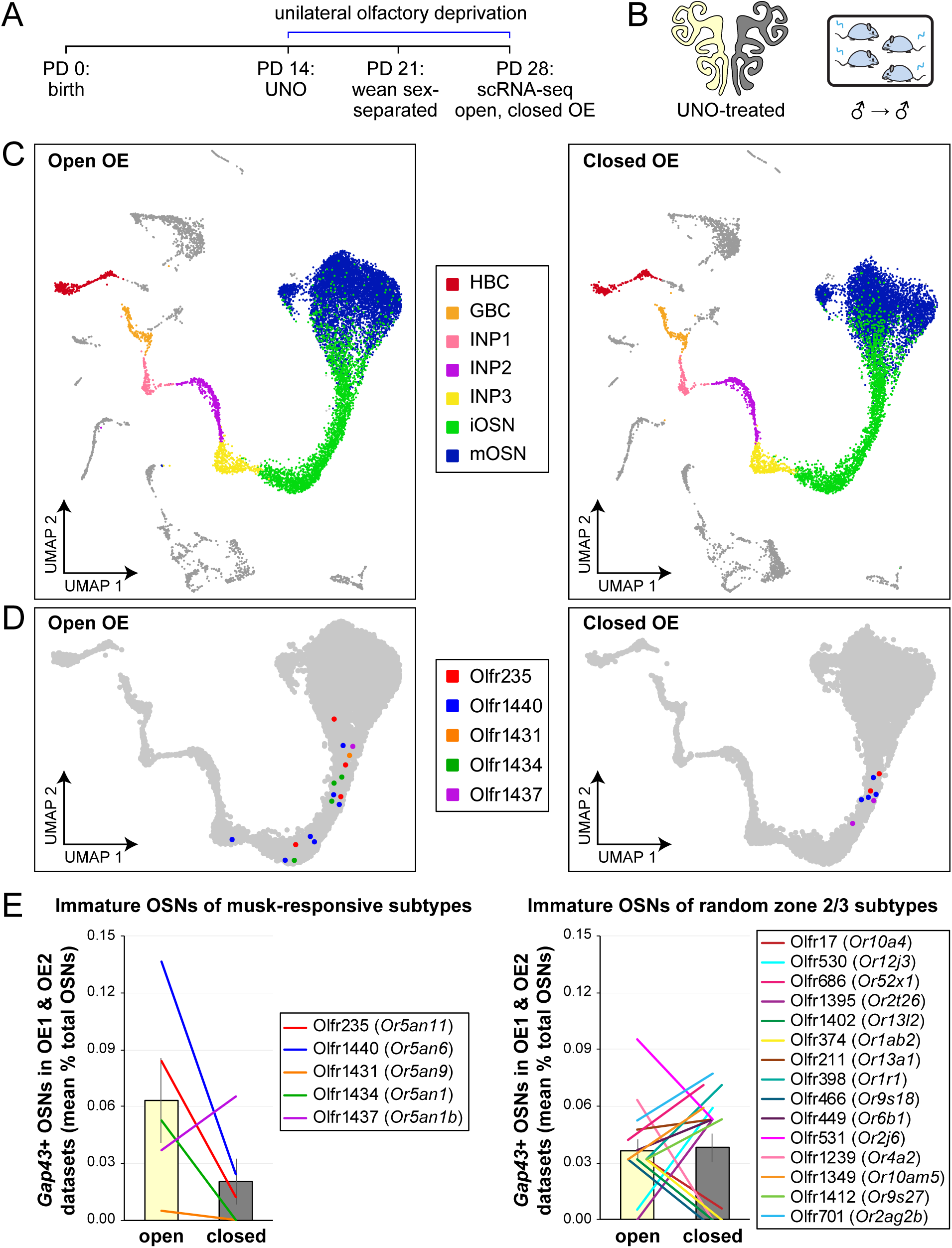
scRNA-seq analyses of the open and closed sides of whole OEs from UNO-treated adolescent male mice reveal greater quantities of newborn OSNs of musk-responsive subtypes on the open side of the OE relative to the closed. A, B. Experimental timeline (A) and conditions (B) used to generate scRNA-seq datasets for assessing the effects of UNO on quantities of newborn OSNs of specific subtypes in the OEs of adolescent male mice. Datasets were generated from the open and closed side of the OEs of mice that were UNO-treated at PD 14, weaned sex-separated at PD 21, and sacrificed at PD 28 ^30^. C. A UMAP representation of all cells within the open (*left*) and closed (*right*) side datasets corresponding to OE sample 2 (OE 2). Cells within the OSN lineage are represented by colored dots, as defined in the legend: horizontal basal cells (HBC; *red*), globose basal cells (GBC; *orange*), immediate neuronal precursor 1 cells (INP1; *pink*), immediate neuronal precursor 2 cells (INP2; *purple*) immediate neuronal precursor 3 cells (INP3; *yellow*), immature OSNs (iOSN; *green*), and mature OSNs (mOSNs; *blue*). D. UMAP representation of cells within the OSN lineage of the open (*left*) and closed (*right*) side datasets of OE 2. Immature (*Gap43*+) OSNs of the 5 known musk-responsive subtypes are represented by colored dots, as indicated in the legend. E. Quantification of individual (*lines*) and mean (*bars*) percentages of the OSN population represented by immature OSNs of musk-responsive subtypes (*left*) or randomly chosen zone 2/3 subtypes (*right*) within the open and closed side datasets. Quantifications represent the averages of subtype-specific iOSN quantities obtained from scRNA-seq datasets corresponding to the OEs from 2 different mice (OE 1 and OE 2). Error bars: SEM. See also Figure 1–figure supplements 1 and 2.

To further test whether olfactory deprivation reduces quantities of newborn OSNs of musk-responsive subtypes in mice exposed to male odors, we employed an established histological assay in which EdU-birthdated OSNs of specific subtypes are identified *via* EdU staining and OR-specific RNA-FISH ^30,46^. Using this approach, we quantified the number of newborn OSNs of four musk-responsive subtypes per half-section, on the open and closed sides of the OEs of male mice that had been UNO-treated at PD 14, weaned sex-separated at PD 21, EdU-injected at PD 28, and dissected at PD 35 (Figure 2A, B). The EdU injection timepoint was based on the timepoint used for identification of immature (*Gap43*-expressing) OSNs in the scRNA-seq datasets, while the dissection timepoint was chosen to provide a chase period of 7 days, which is sufficient to enable robust and stable expression of an OR ^30^. In agreement with findings from scRNA-seq, EdU-labeled OSNs of subtypes Olfr235, Olfr1440, and Olfr1431 exhibited significantly greater quantities on the open side of the OE compared to the closed side in adolescent male mice (*P* < 0.05; 2.3-fold, 1.8-fold, and 2.5-fold, respectively) (Figure 2C-F). By contrast, subtype Olfr1437 did not show a significant open-side bias (*P* > 0.05; 1.01-fold) (Figure 2G), nor did two non-musk-responsive subtypes, Olfr912 and Olfr1463 (*Or5b109*) (1.05- fold for both subtypes) (Figure 2H; Figure 2–figure supplement 1). To verify that observed open-side biases in quantities of newborn OSNs are not an artifact of the normalization method used, which was based on the number of OE half-sections quantified, we compared this method to two alternatives: based on the number of EdU+ cells or on the total DAPI+ area. Notably, all three methods yielded open-side biases of similar magnitude, with no significant differences in UNO effect sizes (defined as the log_2_ [open/closed] ratio) observed between the three methods (Figure 2–figure supplement 2). Taken together, these findings indicate that olfactory deprivation *via* UNO selectively reduces quantities of newborn OSNs of some musk-responsive subtypes in adolescent male mice, consistent with the possibility that the birthrates of these subtypes are selectively accelerated by exposure to male odors.

**Figure 2.**
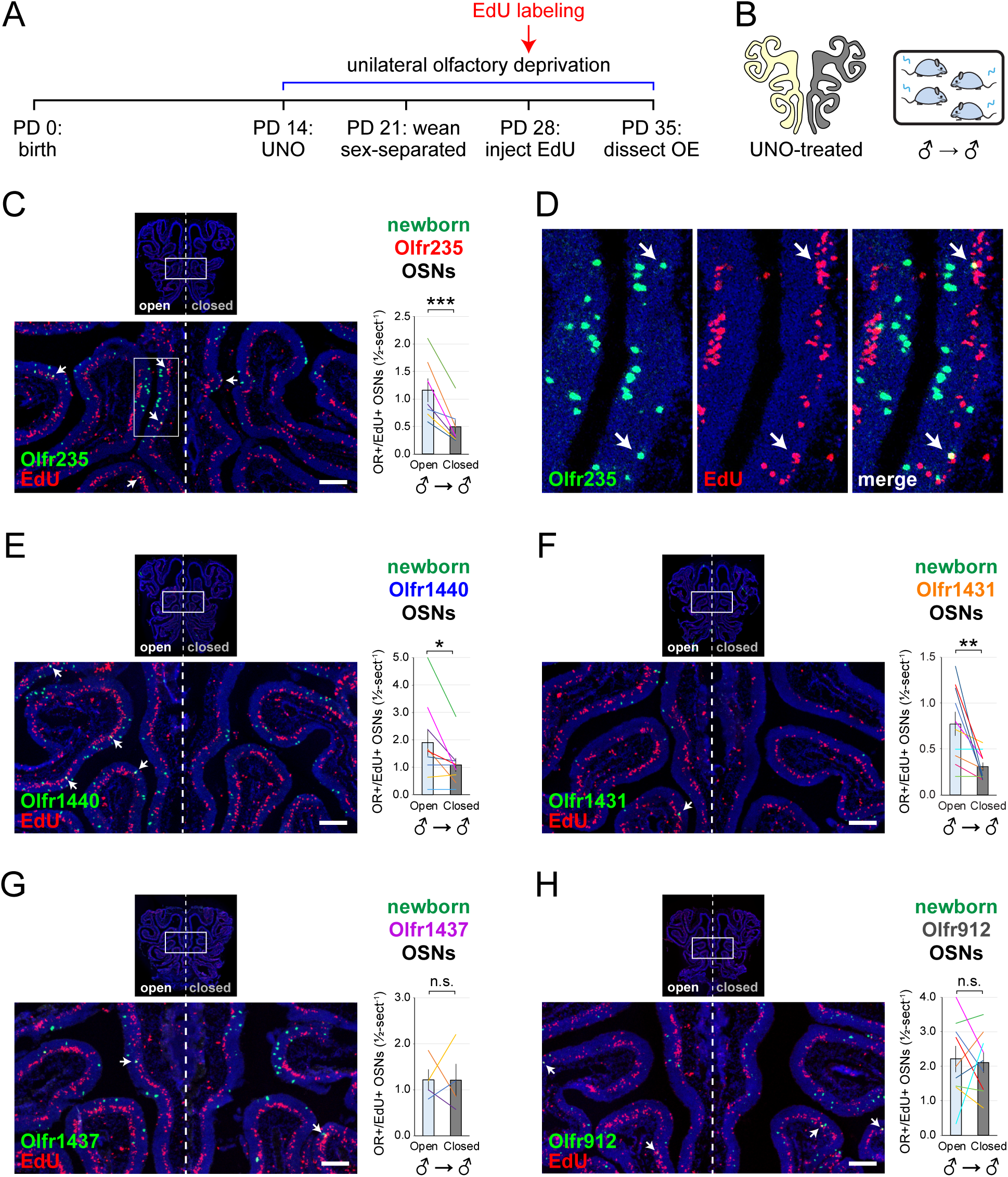
EdU birthdating analyses confirm that UNO-treated adolescent male mice exhibit greater quantities of newborn OSNs of specific musk-responsive subtypes on the open side of the OE relative to the closed. A, B. Experimental timeline (A) and conditions (B) used to generate OE tissue samples for assessing the effects of UNO on quantities of newborn OSNs of specific subtypes in adolescent male mice. Mice were UNO-treated at PD 14, weaned sex-separated at PD 21, EdU-labeled at PD 28, and sacrificed at PD 35. OEs were sectioned and analyzed using OR-specific RNA-FISH and EdU staining. C, E–H. *Left*: Representative images of OE sections from UNO-treated adolescent male mice that were exposed, at the time of EdU labeling, to male littermates (♂ → ♂), with newborn OSNs (OR+/EdU+) indicated by white arrows. *Right*: Quantifications of newborn OSNs on the open and closed sides of tissue sections spanning the anterior-posterior lengths of OEs from UNO-treated male mice reveal significant open-side biases in quantities of newborn OSNs of musk-responsive subtypes Olfr235 (C), Olfr1440 (E), and Olfr1431 (F), but not the musk-responsive subtype Olfr1437 (G) or the SBT-responsive subtype Olfr912 (H). D. Enlarged, split-channel view of boxed region in (C), with Olfr235+/EdU+ OSNs indicated by white arrows. Scale bars: 150 µm. Each line represents a distinct mouse (*n* = 4–10 mice [≥ 5 sections/mouse] per OSN subtype). \*\*\**P* < 0.001; \*\**P* < 0.01; \**P* < 0.05; n.s. *P >* 0.05; ratio paired two-tailed t-test. Error bars: SEM. See also Figure 2– figure supplement 1.

### The exposure of mice to male-specific odors elevates quantities of newborn Olfr235 OSNs

Observations that naris occlusion reduces quantities of newborn OSNs of subtypes Olfr235, Olfr1440, and Olfr1431 in adolescent male mice, together with previous findings that mice exposed to males from weaning until adulthood exhibit elevated representations of all three subtypes, are consistent with the hypothesis that exposure to male odors accelerates the birthrates of these subtypes. If so, we predicted that UNO-induced open-side biases in newborn OSNs of these subtypes may be attenuated in mice that are not exposed to male odors. To test this, we quantified newborn OSNs of subtype Olfr235 in UNO-treated female mice that were housed either sex-separated or sex-combined starting from the age of weaning (PD 21) and thus either unexposed or exposed, respectively, to male odors at the time of EdU labeling (PD 28) (Figure 3A, B; Figure 3–figure supplement 1A). Unlike sex-separated males, sex-separated females showed no significant difference in quantities of newborn Olfr235 OSNs on the open side of the OE compared to the closed (1.1-fold) (Figure 3C, E-*left*) and, correspondingly, a significantly lower UNO effect size (*P* < 0.05; 14-fold) (Figure 3E-*right*). By contrast, sex-combined females showed a significantly greater quantity of newborn Olfr235 OSNs on the open side of the OE compared to the closed (*P* < 0.01; 2.2-fold) and, consequently, a UNO effect size significantly greater than that observed in sex-separated females (*P* < 0.05; 11-fold), but not significantly different from sex-separated males (0.8-fold) (Figure 3D, E). Analogous open-side biases and differences in UNO effect sizes were observed for total Olfr235 OSN quantities (Figure 3–figure supplement 1B; Figure 3F).

**Figure 3.**
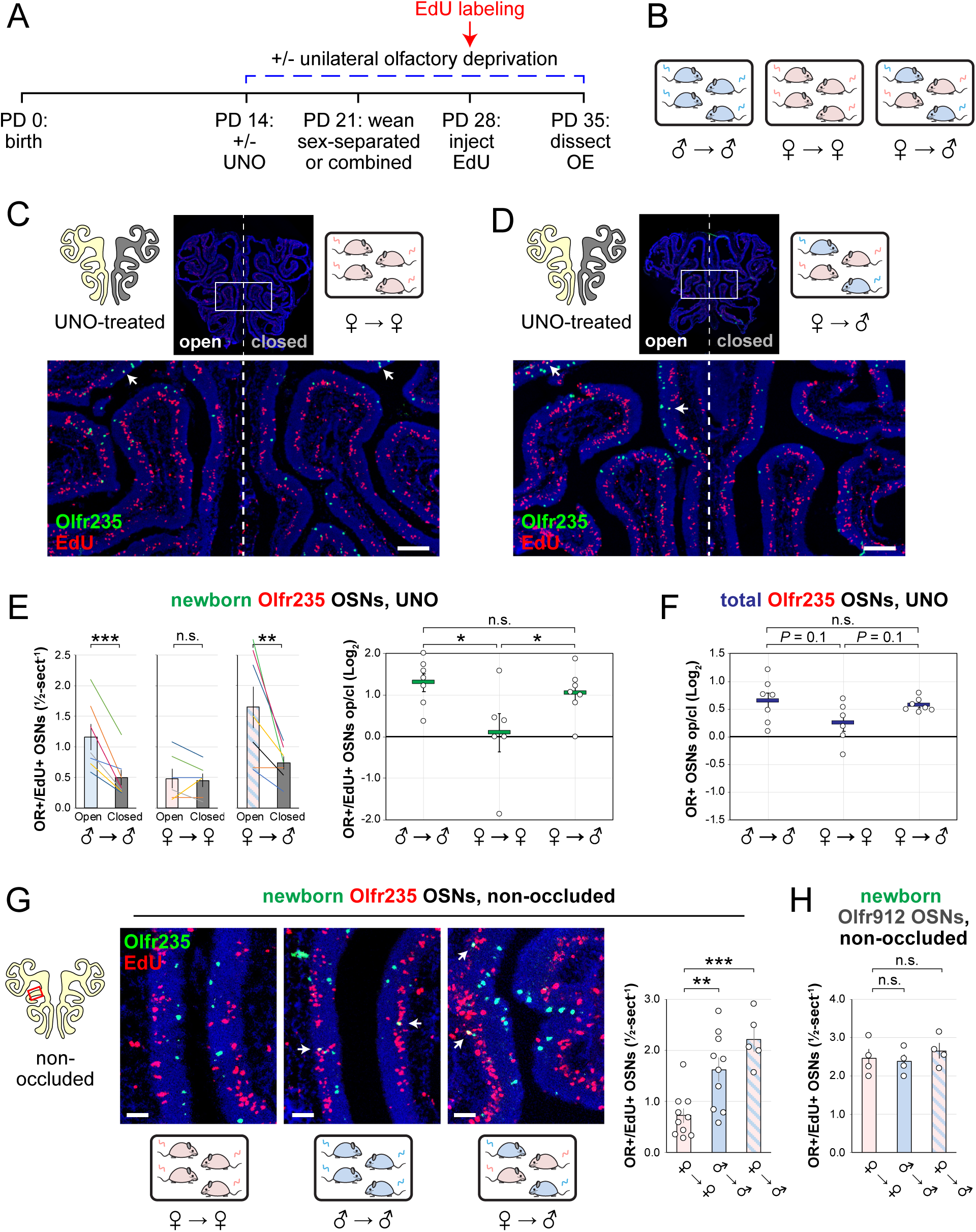
Exposure of mice to male mouse odors causes selective increases in quantities of newborn OSNs of the musk-responsive subtype Olfr235. A, B. Experimental timeline (A) and conditions (B) used to generate OE tissue samples for assessing the effects of exposure to male mouse odors on quantities of newborn OSNs of specific subtypes. Male and female mice were either UNO-treated or untreated (non-occluded) at PD 14, weaned sex-separated or sex-combined at PD 21, EdU-labeled at PD 28, and sacrificed at PD 35. OEs were sectioned and analyzed using OR-specific RNA-FISH and EdU staining. C, D. Representative images of OE sections from UNO-treated female mice that were exposed, at the time of EdU labeling, to either female (♀ → ♀) (C) or male (♀ → ♂) (D) littermates, with newborn Olfr235 OSNs (OR+/EdU+) indicated by white arrows. E, F. Quantifications of newborn (E) and total (F) Olfr235 OSNs on the open and closed sides of tissue sections spanning the anterior-posterior lengths of OEs from UNO-treated mice reveal significant open-side biases in quantities of newborn Olfr235 OSNs in male or female mice exposed to male littermates (♂ → ♂ or ♀ → ♂), but not in females exposed to only female littermates (♀ → ♀) (E-*left*), and greater UNO effect sizes for quantities of newborn (E-*right*) and total (F) Olfr235 OSNs within the OEs of mice that were exposed to male littermates. G-*left, middle*. Representative images (*middle*) corresponding to the region outlined in the schematic (*left, red box*) of OE sections from non-occluded mice that were exposed to male littermates (♂ → ♂, ♀ → ♂) or to female littermates (♀ → ♀) at the time of EdU labeling, with Olfr235+/EdU+ OSNs indicated by white arrows. G-*right*, H. Quantifications of newborn Olfr235 (G-*right*) and Olfr912 (H) OSNs in tissue sections spanning the anterior-posterior lengths of OEs reveal significantly greater quantities of newborn Olfr235 OSNs in non-occluded mice exposed, at the time of EdU labeling, to male littermates (♂ → ♂, ♀ → ♂) compared to those exposed only to female littermates (♀ → ♀), while quantities of newborn Olfr912 OSNs showed no significant differences between mice exposed to male littermates and those exposed only to females. Scale bars: 150 µm (C, D), 50 µm (G). Each line or circle represents a distinct mouse (*n* = 4–10 mice [≥ 5 sections/mouse] per OSN subtype and condition). \*\*\**P* < 0.001; \*\**P* < 0.01; \**P* < 0.05; n.s. *P>* 0.05; ratio paired two-tailed t-test (E, *left*); one-way ANOVA test, FDR-adjusted (E, *right*, F-H). Error bars: SEM. Data in panel E for newborn OSN quantities in ♂ → ♂ OE samples correspond to Figure 2. See also Figure 3–figure supplements 1-3.

Findings that exposure to male odors is required for occlusion-induced reductions in quantities of newborn Olfr235 OSNs are consistent with the hypothesis that a component of male odors can accelerate the birthrate of this subtype. If so, the exposure of non-occluded mice to male odors should increase observed quantities of newborn Olfr235 OSNs. To test this, we compared quantities of newborn Olfr235 OSNs in non-occluded sex-separated females, sex-separated males, and sex-combined females (Figure 3A, B). Remarkably, compared to sex-separated females, sex-separated males and sex-combined females exhibited significantly greater quantities of newborn Olfr235 OSNs (*P* < 0.01; 2.2-fold and 3.1-fold, respectively) (Figure 3G). By contrast, quantities of newborn OSNs of the SBT-responsive subtype Olfr912 showed no significant differences between the three experimental groups (Figure 3H). These findings strongly support the conclusion that the exposure of mice to male odors increases quantities of newborn Olfr235 OSNs within the OE.

Unlike subtype Olfr235, subtypes Olfr1440 and Olfr1431 exhibited significant open-side biases in both newborn and total OSN quantities within the OEs of UNO-treated sex-separated males, sex-separated females, and sex-combined females (*P* < 0.05), with no significant differences in UNO effect size observed between the three groups (Figure 3–figure supplement 1C-F). As expected, the non-musk-responsive OSN subtypes Olfr912 and Olfr1463 exhibited no significant open-side biases in newborn OSN quantities in any experimental group, although Olfr912 showed significant closed-side biases in total OSNs (*P* < 0.01) (Figure 3–figure supplement 1G-J), consistent with the possibility that naris occlusion protects OSNs of this subtype from overstimulation ^19,24^. Observations that exposure to male odors is required for occlusion-induced reductions in quantities of newborn OSNs of subtypes Olfr235 but not Olfr1440 or Olfr1431 were unexpected considering that ORs of all three subtypes show male-biased expression in mice housed sex-separated until 6 months of age (Appendix 1–figure 1) ^19^. A conceivable explanation for these findings is that, unlike subtype Olfr235, quantities of newborn Olfr1440 and Olfr1431 OSNs are elevated by exposure to odors emitted by both male and female mice at the adolescent stage. Such differences could reflect variations in the specific odorants to which distinct musk-responsive subtypes are most sensitive ^33^, and which may depend on the age and sex of mice contributing to the odor environment ^47–50^. If so, we predicted that the age and/or sex of mice within the odor environment at the time of EdU labeling might differentially affect the degree to which quantities of newborn OSNs of these subtypes are affected by olfactory deprivation. To test this, we compared open-side biases in newborn OSN quantities within UNO-treated adolescent males exposed to littermates alone versus those exposed to both littermates and adult parents (Figure 3–figure supplement 3A, B). Remarkably, newborn Olfr1431 OSNs exhibited a significantly greater (*P* < 0.05; 2-fold) UNO effect size in the presence of adult mice compared to the absence, reflecting open-side biases of 4.8-fold and 2.5-fold, respectively (Figure 3–figure supplement 3C, D). By contrast, UNO effect sizes for newborn OSNs of subtypes Olfr235, Olfr1440, and Olfr1437, as well as the control subtype Olfr912, were not significantly affected by exposure to adults (Figure 3–figure supplement 3E-I). These findings are consistent with the possibility that stimulation-dependent changes in the quantities of newborn OSNs of musk-responsive subtypes may vary depending on the age of odor-emitting mice within the environment.

### The exposure of UNO-treated mice to low concentrations of a musk odorant increases open-side biases in quantities of newborn OSNs of musk-responsive subtypes

Evidence that quantities of newborn OSNs of musk-responsive subtypes can be affected by exposure to unknown components of mouse odors indicated that the birthrates of specific OSN subtypes may be accelerated by discrete odorants. If so, we predicted that the exposure of mice to musk odorants that are known to stimulate these subtypes would increase their neurogenesis rates. However, because high levels of chronic stimulation of OSNs have been found to reduce the representations of stimulated subtypes within the OSN population, presumably *via* reductions in OSN lifespan ^12,13,19^, we speculated that there may be a range of odorant concentrations sufficient to accelerate the birthrates of these subtypes without shortening their OSN lifespans. To assess this, we analyzed the effects of exposing UNO-treated female mice to varying concentrations of the musk odorant (*R*)-3-methylcyclopentadecanone (muscone) on quantities of newborn and total OSNs of the musk-responsive subtypes Olfr235, Olfr1440, and Olfr1431, as well as the control subtypes Olfr912 and Olfr1463, *via* a metal tea-ball from weaning (PD 21) until dissection (PD 35) (Figure 4A, B). Remarkably, exposure of mice to muscone was found to significantly affect the extent to which UNO treatment alters the relative quantities of both newborn and total OSNs of musk-responsive subtypes on the open and closed sides of the OE in a concentration-dependent manner (Figure 4C; Figure 4–figure supplements 1, 2). Subtype Olfr235, for example, which showed no significant difference in newborn OSN quantities between the open and closed sides in unexposed females, exhibited significant open-side biases in females exposed to 0.1% or 1% muscone (*P* < 0.01; 2.4 and 2.0-fold, respectively), but not those exposed to 10% (1.2-fold) (Figure 4C-*middle*). Accordingly, mice exposed to 0.1% or 1% muscone exhibited significantly greater UNO effect sizes for quantities of newborn Olfr235 OSNs compared to unexposed females (*P* < 0.05; 14 and 13-fold, respectively) (Figure 4C- *right*). Similarly, open-side biases in total Olfr235 OSNs were observed in mice exposed to 0.1% or 1% muscone, with the most significant effect observed at 0.1% (*P* < 0.01) (Figure 4–figure supplement 2). One possible explanation for the reduction in UNO effect sizes observed at 10% muscone relative to lower concentrations is that small amounts of odorant may enter the closed side of the OE due to transnasal and/or retronasal odor transfer *via* the nasopharyngeal canal ^51^ and nasopharynx, respectively ^31^, an effect that would be expected to increase with environmental concentrations and attenuate differences in observed effects of odor stimulation between the open and closed sides.

**Figure 4.**
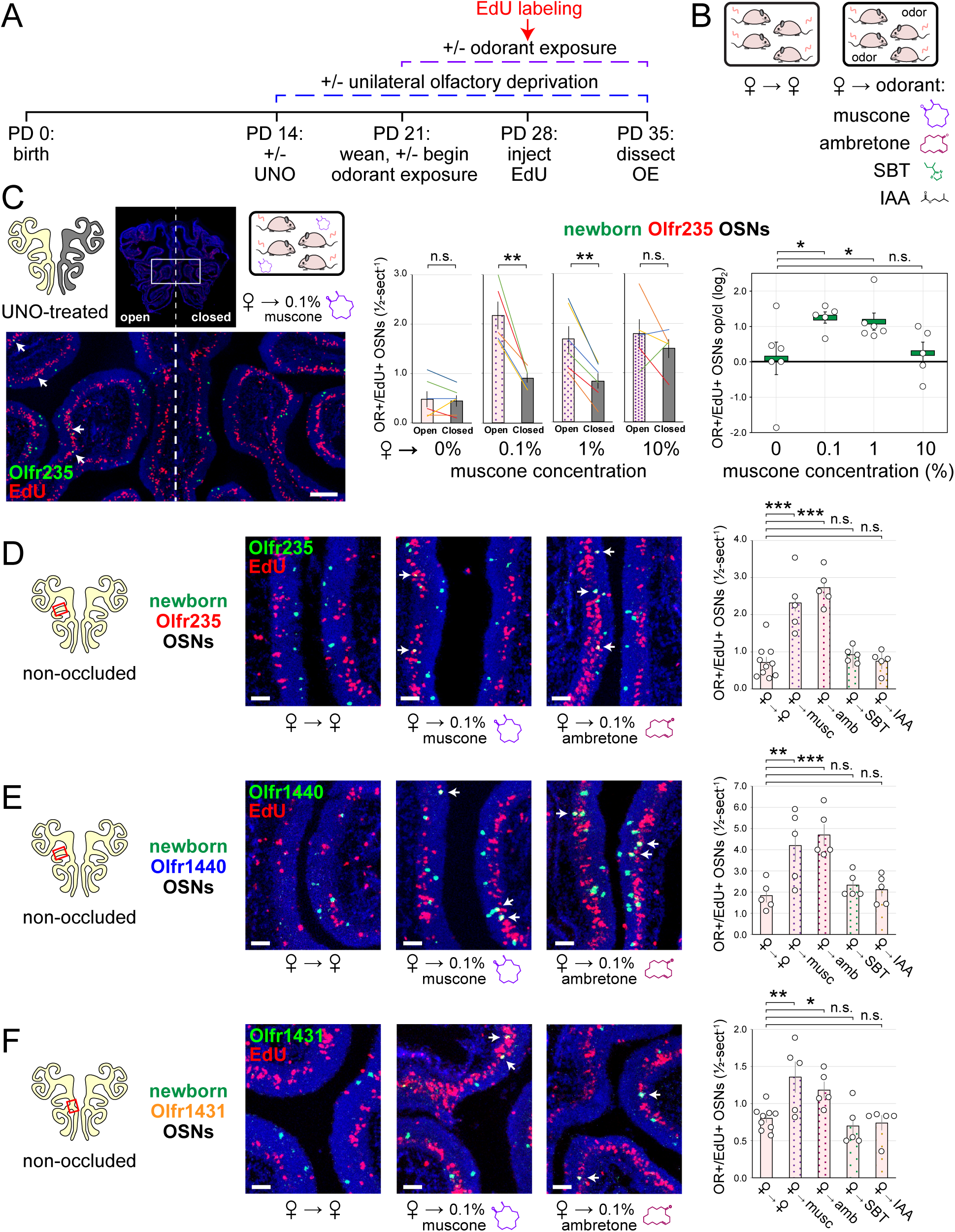
Exposure of mice to musk odors causes selective increases in quantities of newborn OSNs of musk-responsive subtypes. A, B. Experimental timeline (A) and conditions (B) used to generate OE tissue samples for assessing the effects of exposure to musk (muscone, ambretone) or non-musk (SBT, IAA) odorants on quantities of newborn OSNs of specific subtypes. Female mice were either UNO-treated or untreated (non-occluded) at PD 14, either exposed or unexposed to an exogenous musk or non-musk odorant starting at PD 21, EdU-labeled at PD 28, and sacrificed at PD 35. OEs were sectioned and analyzed using OR-specific RNA-FISH and EdU staining. C. *Left*: Representative image of an OE section from a UNO-treated female mouse that was exposed, at the time of EdU labeling, to 0.1% muscone, with newborn Olfr235 OSNs (OR+/EdU+) indicated by white arrows. *Middle, right*: Quantifications of newborn Olfr235 OSNs on the open and closed sides of tissue sections spanning the anterior-posterior lengths of OEs from UNO-treated female mice reveal significant open-side biases (*middle*) and significantly greater UNO effect sizes (*right*) in mice exposed to 0.1% or 1% (but not 10%) muscone compared to 0%, with a maximum effect size observed at a concentration of 0.1%. D–F. *Left, middle*: Representative images (*middle*) corresponding to the regions outlined in schematics (*left, red box*) of OE sections from non-occluded female mice that were exposed just to female littermates (♀ → ♀) or also to 0.1% muscone or 0.1% ambretone, with newborn Olfr235 (D), Olfr1440 (E), and Olfr1431 (F) OSNs (OR+/EdU+) indicated by white arrows. *Right*: Quantifications of newborn Olfr235 (D), Olfr1440 (E), and Olfr1431 (F) OSNs in tissue sections spanning the anterior-posterior lengths of OEs reveal that, compared to females exposed just to female littermates (♀ → ♀), significantly greater quantities of newborn OSNs of all 3 subtypes were observed in females also exposed to muscone (musc; 0.1%) or ambretone (amb; 0.1%), but not the non-musk odorants SBT (0.1%) or IAA (0.1%). Scale bars: 150 µm (C), 50 µm (D–F). Each line or circle represents a distinct mouse (*n* = 5–10 mice [≥ 5 OE sections/mouse] per OSN subtype and condition). \*\*\**P* < 0.001; \*\**P* < 0.01; \**P* < 0.05; n.s. *P>* 0.05; ratio paired two-tailed t-test (C, *middle*); one-way ANOVA test, FDR-adjusted (C, *right*, D–F). Error bars: SEM. Data for ♀ → 0% muscone samples in panel C correspond to Figure 3E. Image and data for ♀ → ♀ samples (D) correspond to Figure 3G. See also Figure 4–figure supplements 1–4.

Compared to subtype Olfr235, open-side biases in quantities of newborn and total OSNs of subtypes Olfr1440 and Olfr1431 were more subtly affected by the concentration of muscone to which mice were exposed (Figure 4–figure supplements 1, 2). We speculate that this may be explained in part by the fact that both subtypes exhibit significant open-side biases in UNO- treated female mice even in the absence of exogenous odor exposure, which may limit the influence of muscone exposure on the UNO effect size. In the case of Olfr1440, the subtype most sensitive to muscone ^33,34^, it is conceivable that mature OSNs are robustly stimulated on the closed side of the OE by very small amounts of odors that enter transnasally and/or retronasally. Consistent with this possibility, UNO-treated females exposed to 0.1% muscone show greater quantities of newborn Olfr1440 OSNs on both the open and closed sides of the OE compared to their unexposed counterparts (Figure 4–figure supplement 1A-middle). Relatedly, subtype Olfr1440 showed evidence of shortened OSN lifespans in mice exposed to 1% and 10% muscone, as reflected in the higher total OSN quantities observed on the closed side of the OE (Figure 4–figure supplement 2B). In the case of Olfr1431, newborn OSNs exhibited a significantly greater open-side bias in mice exposed to 1% muscone compared to unexposed controls (*P* < 0.01; 1.6-fold), but no significant difference at 0.1%, perhaps indicating that this subtype is less sensitive to muscone than other subtypes. Like subtype Olfr235, neither Olfr1440 nor Olfr1431 exhibited significant open-side biases in newborn OSN quantities in mice exposed to 10% muscone, potentially reflecting substantial transnasal and/or retronasal odor transfer to the closed side of the OE at this concentration. As expected, muscone exposure did not significantly affect open-side biases in newborn or total OSN quantities of the non-musk responsive subtypes Olfr912 or Olfr1463 (Figure 4–figure supplements 1, 2). Taken together, these findings reveal that the exposure of UNO-treated mice to low concentrations of a musk odorant is sufficient to induce open-side biases in the quantities of newborn OSNs of musk-responsive subtypes without apparent adverse effects on OSN survival.

### The exposure of non-occluded mice to musk odorants selectively elevates quantities of newborn OSNs of musk-responsive subtypes

Findings that the exposure of UNO-treated mice to muscone concentrations as low as 0.1% can increase open-side biases in quantities of newborn OSNs of musk-responsive subtypes suggested that the exposure of non-occluded mice to musk odorants at this concentration might selectively accelerate the birthrates of these subtypes. To test this, we compared quantities of newborn OSNs of three musk-responsive subtypes and two musk non-responsive subtypes within the OEs of non-occluded female mice exposed to a 0.1% concentration of one of two different musk odorants, muscone or 5-Cyclohexadecenone (ambretone), one of two different non-musk odorants, SBT or isoamyl acetate (IAA), or no odorant. Relative to unexposed controls, all three musk-responsive subtypes showed significantly elevated quantities of newborn OSNs in mice exposed to muscone and ambretone (*P* < 0.05), with Olfr235 exhibiting increases of 3.2 and 3.8- fold, Olfr1440 exhibiting increases of 2.3 and 2.6-fold, and Olfr1431 exhibiting increases of 1.7 and 1.5-fold, respectively (Figure 4D-F). Notably, these differences were not affected by the use of alternative normalization methods (Figure 4–figure supplement 3). By contrast, none of the three musk-responsive subtypes exhibited significant changes in newborn OSN quantities in mice exposed to the non-musk odorants SBT or IAA. Likewise, the SBT-responsive subtype Olfr912 exhibited no significant changes in quantities of newborn OSNs in mice exposed to muscone, ambretone, or SBT (Figure 4–figure supplement 4C-*left*). Moreover, two non-musk-responsive subtypes that were previously found to undergo occlusion-dependent reductions in neurogenesis ^30^, Olfr827 (*Or9k7*) and Olfr1325 (*Or13ae2*), showed no significant changes in quantities of newborn OSNs in mice exposed to muscone (Figure 4–figure supplement 4C-*middle, right*). Taken together, these findings indicate that: 1) the exposure of non-occluded mice to exogenous musk odorants selectively increases quantities of newborn OSNs of musk-responsive subtypes, 2) the exposure of mice to exogenous non-musk odorants has no effect on quantities of newborn OSNs of musk-responsive subtypes, and 3) only a fraction of subtypes have a capacity to exhibit increases in quantities of newborn OSNs following exposure to their cognate odorants.

### Musk odorant-dependent increases in quantities of newborn OSNs of musk-responsive subtypes persist into adulthood

Findings that the exposure of adolescent mice to male or musk-like odors elevates quantities of newborn OSNs of musk-responsive subtypes raise the question of whether this phenomenon is limited to early life or, rather, persists into adulthood. To address this, we compared quantities of newborn OSNs of three musk-responsive subtypes and one musk non-responsive subtype within the OEs of 9-week-old (PD 65) non-occluded female mice that had been either unexposed to an exogenous odorant or exposed to 0.1% muscone starting from weaning (PD 21) and EdU-treated for 3 days starting at 8 weeks of age (PD 56-58) (Figure 5A, B). Remarkably, despite the generally reduced OSN birthrate in adult mice compared to adolescents, all three musk-responsive subtypes showed elevated quantities of newborn OSNs in mice exposed to muscone relative to unexposed controls, with Olfr235, Olfr1440 and Olfr1431 exhibiting 1.5-fold, 2.2- fold, and 1.9-fold increases, respectively (Figure 5C-E). Notably, observed increases reached statistical significance (*P* < 0.05) only for subtype Olfr1440, likely due to the higher baseline birthrate of this subtype relative to Olfr235 and Olfr1431. As expected, exposure to muscone did not significantly increase quantities of newborn OSNs of subtype Olfr912 (Figure 5F). These findings indicate that the capacity for odorant-dependent increases in quantities of newborn OSNs of specific subtypes persists into adulthood.

**Figure 5.**
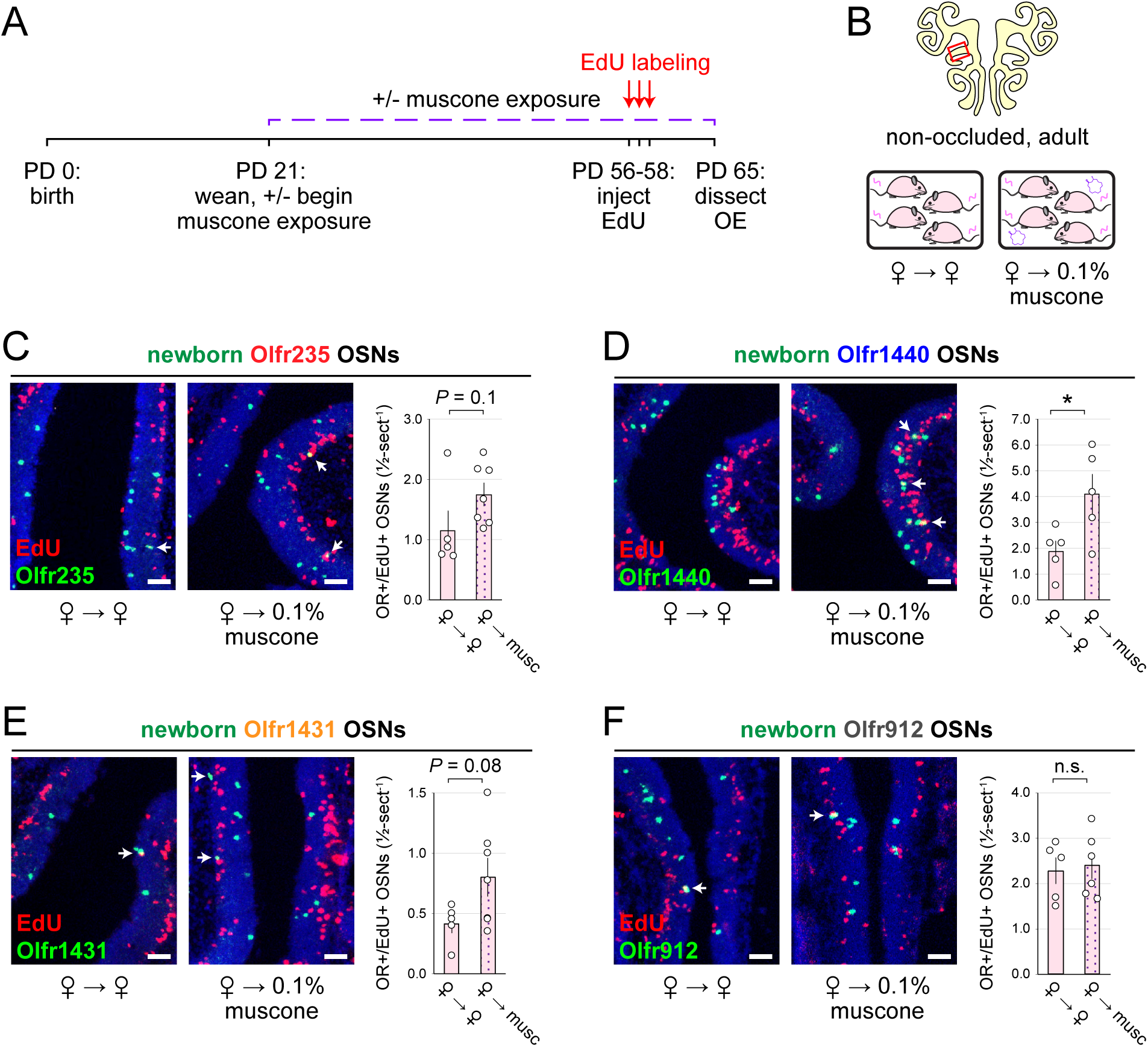
Musk exposure increases quantities of newborn OSNs of musk-responsive subtypes in adulthood. A, B. Experimental timeline (A) and conditions (B) used to generate OE tissue samples for assessing the effects of exposure to musk odors on quantities of newborn OSNs of specific subtypes in non-occluded adult mice. Mice were either exposed or unexposed to 0.1% muscone starting at PD 21, EdU-labeled at PD 56-58, and sacrificed at PD 65. OEs were sectioned and analyzed using OR-specific RNA-FISH and EdU staining. C–F. *Left*: Representative images corresponding to the region outlined in the schematic in panel B (*top*), of OEs from non-occluded adult female mice that were either exposed to just female littermates (♀ → ♀) or also to muscone (♀ → 0.1% muscone), with newborn Olfr235 (C), Olfr1440 (D), Olfr1431 (E), and Olfr912 (F) OSNs (OR+/EdU+) indicated by white arrows. *Right*: Quantifications of newborn OSNs within tissue sections spanning the anterior-posterior lengths of OE from non-occluded adult female mice reveal greater quantities of newborn OSNs of all 3 musk-responsive subtypes (Olfr235, Olfr1440, and Olfr1431) in mice exposed to muscone compared to just female littermates (♀ → ♀), while quantities of newborn OSNs of the SBT- responsive subtype Olfr912 were relatively unaffected. Scale bars: 50 µm. Each circle represents a distinct mouse (*n* = 5–7 mice [≥ 5 OE sections/mouse] per OSN subtype and condition). \**P* < 0.05; n.s. *P>* 0.05; unpaired two-tailed *t*-test. Error bars: SEM.

### The time-independence of odor-driven increases in EdU-labeled OSNs of musk-responsive subtypes is consistent with a mechanism involving altered OSN birthrates

Odor-dependent increases in the quantities of newborn OSNs of musk-responsive subtypes could, in principle, be caused by a mechanism that selectively accelerates the rates with which these subtypes are generated or, alternatively, the rates with which they are selectively enriched following their generation (e.g., *via* enhanced survival or OR switching ^52^). If differences in newborn OSN quantities are mediated by selective enrichment, they would be expected to increase in magnitude over time following EdU labeling, as a subset of newborn OSNs exhibit longer lifespans or switch their OR identities in the presence of stimulation. If, however, changes are mediated by accelerated birthrates of specific OSN subtypes, increases in newborn OSN quantities should appear shortly following EdU labeling and remain stable over time. To distinguish between these possibilities, we compared stimulation-dependent changes in quantities of newborn musk-responsive OSNs at two timepoints: 4 days post-EdU, the earliest point during OSN differentiation when OR transcripts can be consistently detected *via* RNA-FISH ^30,53^, and three days later (7 days post-EdU) (Figure 6A). In initial experiments, the time-dependence of open-side biases in quantities of EdU-labeled OSNs of musk-responsive subtypes were assessed in UNO-treated and sex-separated males, females, and females exposed to 0.1% muscone (Figure 6B). In sex-separated males, robust open-side biases and statistically indistinguishable UNO effect sizes were observed at 4 and 7 days-post EdU for quantities of EdU-labeled OSNs of subtypes Olfr235, Olfr1440, and Olfr1431 (Figure 6C, D; Figure 6–figure supplement 1A, B). Likewise, UNO-treated and muscone-exposed females exhibited robust open-side biases in quantities of newborn Olfr235 OSNs and statistically indistinguishable UNO effect sizes at 4 and 7 days-post EdU (Figure 6E; Figure 6–figure supplement 1C). As expected, no significant open-side biases in EdU-labeled OSN quantities or UNO effect sizes were observed at either 4 or 7 days post-EdU for subtypes Olfr912 or Olfr1463 in UNO-treated male (Figure 6–figure supplement 1D, E) or female mice exposed to muscone (Figure 6–figure supplement 1F), or for subtype Olfr235 in sex-separated females (Figure 6–figure supplement 1G).

**Figure 6.**
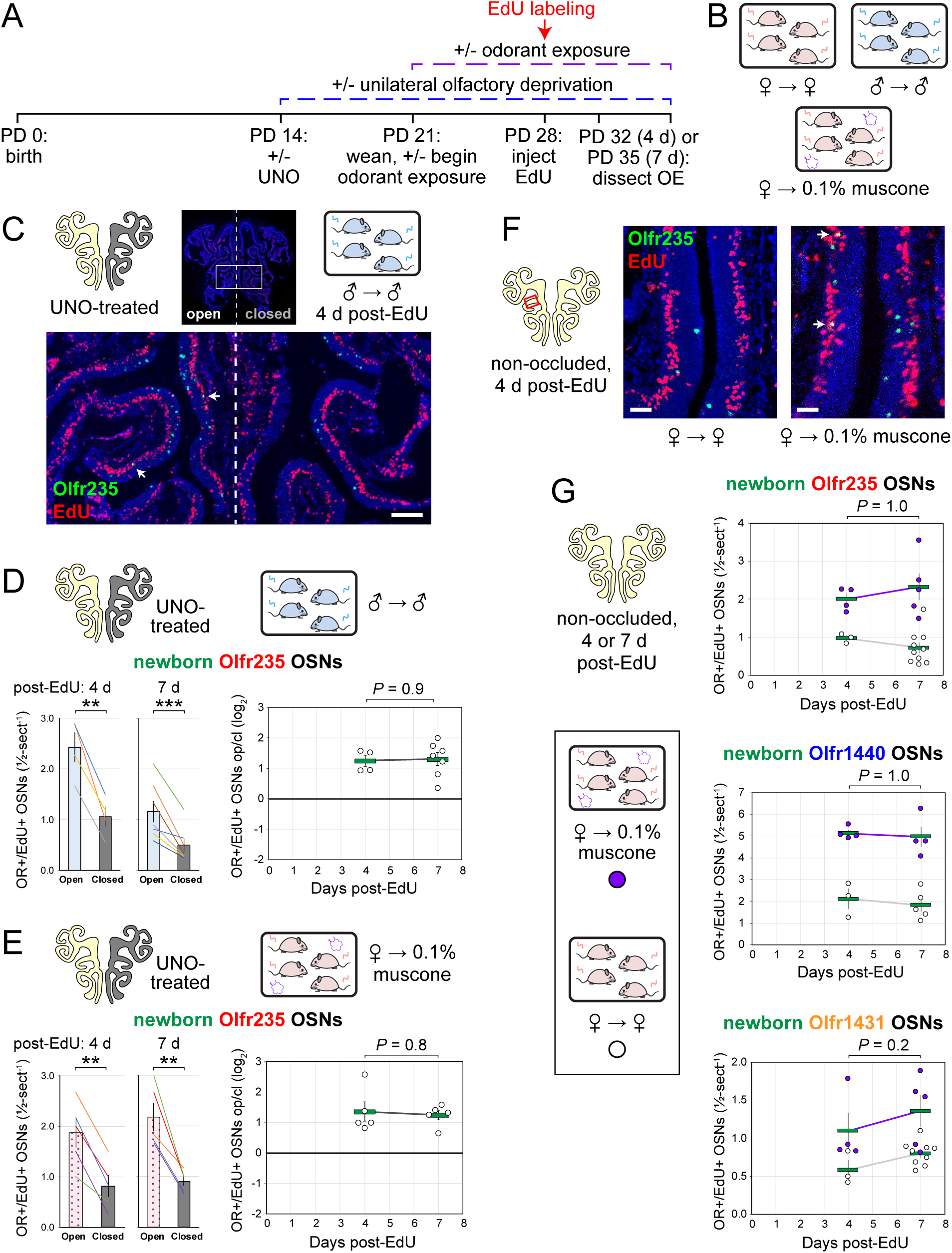
Stimulation-dependent increases in quantities of newborn OSNs of musk-responsive subtypes are stable over time following birth, consistent with a mechanism involving altered neurogenesis. A, B. Experimental timeline (A) and conditions (B) used to generate OE tissue samples for assessing the time-dependence of the effects of exposure to male or musk odors on quantities of newborn OSNs of specific subtypes. Mice were either UNO-treated or untreated (non-occluded) at PD 14, either exposed or unexposed to muscone starting at PD 21, EdU- labeled at PD 28, and sacrificed at either PD 32 (4 d post-EdU) or PD 35 (7 d post-EdU). OEs were sectioned and analyzed using OR-specific RNA-FISH and EdU staining. C. Representative image of an OE section from a UNO-treated male mouse that was exposed, at the time of EdU labeling, to male littermates (♂ → ♂) and sacrificed 4 d post-EdU, with newborn Olfr235 OSNs (OR+/EdU+) indicated by white arrows. D, E. Quantifications of newborn Olfr235 OSNs on the open and closed sides of tissue sections spanning the anterior-posterior lengths of OEs from UNO-treated male mice that were exposed to male littermates (♂ → ♂) (D) or female mice that were exposed to 0.1% muscone (♀ → 0.1% muscone) (E) and sacrificed 4 or 7 d post-EdU. Under both conditions, significant open-side biases in quantities of newborn Olfr235 OSNs were observed at both timepoints (*left*), with no statistically significant differences in UNO effect sizes observed over time (*right*). F. Representative images (*right*) corresponding to the region outlined in schematic (*left, red box*) of OE sections from non-occluded female mice that were exposed, at the time of EdU labeling, to either just their female littermates (♀ → ♀) or also to muscone (♀ → 0.1% muscone) and sacrificed 4 d post-EdU, with newborn Olfr235 OSNs (OR+/EdU+) indicated by white arrows. G. Quantifications of newborn Olfr235 (*top*), Olfr1440 (*middle*), and Olfr1431 (*bottom*) OSNs within tissue sections spanning the anterior-posterior lengths of OEs from non-occluded female mice that exposed to either just their female littermates (♀ → ♀) or also to muscone (♀ → 0.1% muscone) and sacrificed 4 or 7 d post-EdU. Greater quantities of newborn OSNs of all 3 subtypes were observed at both timepoints in muscone-exposed mice compared to controls, with no statistically significant interaction between time and newborn OSN quantities observed. Scale bars: 150 µm (C), 50 µm (F). Each circle represents a distinct mouse (*n* = 4–10 mice [≥ 5 OE sections/mouse] per OSN subtype and condition). \*\*\**P* < 0.001; \*\**P* < 0.01; ratio paired two-tailed t-test (D, E *left*); unpaired two-tailed t-test (D, E *right*); two-way ANOVA test (factors: muscone exposure, days post-EdU) (G). Error bars: SEM. Data for 7 d post-EdU samples correspond to Figures 2–4. See also Figure 6–figure supplement 1.

Using a similar approach, we also assessed the time-dependence of differences in muscone-driven increases in quantities of EdU-labeled OSNs of specific subtypes in non-occluded female mice (Figure 6G-*left*). Consistent with UNO-based findings, non-occluded mice exhibited statistically indistinguishable muscone-dependent increases at 4 and 7 days post-EdU for quantities of newborn OSNs of the musk-responsive subtypes Olfr235, Olfr1440, and Olfr1431 (Figure 6G-*right*), as well as the control subtype Olfr912 (Figure 6–figure supplement 1H). Taken together, these findings support the hypothesis that odor-dependent increases in the quantities of newborn OSNs of musk responsive subtypes reflect selective changes in the birthrates of these subtypes.

### Evidence that mice emit musk-like odors

Previous findings that OSNs of musk-responsive subtypes respond to male mouse odors ^19^ and become more highly represented within the OEs of mice exposed to males ^19,24^, together with evidence from the current study that the exposure of mice to male odors can accelerate the birthrates of these subtypes, indicate that mice emit odors that stimulate musk-responsive subtypes. Interestingly, mouse preputial glands have been found to have a high degree of histomorphological, transcriptomic, and molecular similarities to muskrat scent glands ^54^, which secrete musk odors ^55^. Moreover, a previous study noted preliminary evidence that mouse preputial gland extracts contain compounds that activate Olfr1440 OSNs ^33^. Based on this information, we analyzed mouse preputial gland extracts *via* gas chromatography – mass spectrometry (GC-MS) for known musk molecules, particularly those shown to stimulate Olfr235 and Olfr1440 OSNs ^33^. Intriguingly, these analyses revealed the presence of molecules whose GC-MS signals are structurally consistent with known musk molecules (Appendix 2– figure 1). A potential match to one such signal is cycloheptadecanol, a musk molecule that is structurally similar to those known to activate Olfr235 and Olfr1440 OSNs ^33^. Verification of the presence of this and other musk-like odors that correspond to signals from mouse preputial glands will require comparisons to pure standards. Notably, the approach employed here was solely focused on the preputial gland and known musk molecules. However, it is conceivable that mouse musk-responsive OSNs are naturally stimulated by molecules that are structurally unrelated to those that have been characterized to date or are emitted from sources other than preputial glands. Future studies will be needed to address these possibilities.

## Discussion

### The exposure of mice to discrete odorants can selectively accelerate the birthrates of OSN subtypes that they stimulate

Previous studies have established that UNO reduces the overall rate of neurogenesis on the closed side of the OE relative to the open ^27–29,56^. Recently, these occlusion-induced reductions were found to reflect decelerations of the birthrates of only a fraction of the ∼1200 OSN subtypes and to vary according to age and/or the olfactory environment ^30^, suggesting the possibility that unknown olfactory stimuli selectively promote the birthrates of these subtypes. In this study we aimed to test this by elucidating the nature of stimuli that accelerate the birthrates of specific OSN subtypes. We have presented findings that these stimuli include discrete odorants that selectively activate the same subtypes whose birthrates are accelerated. These findings build upon previous observations that a small group of musk-responsive OSN subtypes are more highly represented in the OEs of mice exposed to male odors compared to mice isolated from them ^19,24^, and that some of these subtypes are also responsive to male-specific odors ^19^ (Appendix 1–figure 1). These data suggested that one or more components of mouse-emitted odors might naturally accelerate the birthrates of these OSN subtypes. Here, using scRNA-seq-based and histological approaches, we have found that UNO treatment of adolescent male mice reduces quantities of newborn OSNs of musk-responsive subtypes on the closed side of the OE relative to the open, consistent with the possibility that these subtypes have a capacity to undergo stimulation-dependent neurogenesis (Figures 1, 2). Additionally, we have presented findings that the exposure of mice to male-specific odors or two different exogenous musk odorants can selectively elevate quantities of newborn OSNs of musk-responsive subtypes (Figures 3-5). Finally, we have described evidence that odor-driven increases in newborn OSN quantities are stable over time after EdU labeling, indicating that the observed changes reflect altered OSN birthrates as opposed to altered OSN lifespan or OR gene switching (Figure 6). Collectively, these findings support the hypothesis that the stimuli that regulate the birthrates of specific subtypes are discrete odors that selectively stimulate those subtypes.

### How do discrete odors accelerate the birthrates of specific OSN subtypes?

Our findings that specific OSN subtypes exhibit accelerated birthrates following the exposure of mice to odors that selectively stimulate OSNs of these subtypes indicate that this process occurs *via* a mechanism that is highly specific with respect to the stimulating odors and the subtypes whose birthrates are accelerated. Considering that horizontal basal cells (HBCs) and globose basal cells (GBCs), the stem and progenitor cells that give rise to new OSNs, lack ORs and signal transduction molecules needed to detect and respond to odors, we hypothesize the existence of a signaling pathway from mature OSNs to HBCs or GBCs that alters the rates at which OSNs of specific subtypes are born. Findings from the present study and a previous one ^30^ indicate that this signaling capacity may be limited to only a fraction of OSN subtypes, since a majority of subtypes do not appear to exhibit stimulation-dependent neurogenesis ^30^. Subtype Olfr912, for example, which detects the male-specific odor component SBT ^24^, exhibits no increase in birthrate upon exposure of female mice to male odors, and was therefore employed in this study as a control subtype. We speculate that the receipt of odor-derived signals by HBCs or GBCs alters OR choices or amplifies choices that have already been made. Elucidating the nature of odor-dependent signals received by HBCs/GBCs, as well as the mechanism by which these signals accelerate the birthrates of specific OSN subtypes are important areas of future investigation.

### What functions does discrete odor stimulation-dependent neurogenesis of specific OSN subtypes serve?

Because OSN differentiation entails the stochastic process of singular OR choice ^9,10^, it has long been assumed that OSN neurogenesis is entirely stochastic with respect to OR identity. Thus, unlike other regions of the nervous system where persistent neurogenesis is known to play important adaptive roles ^6–8^, life-long neurogenesis within the OE is generally assumed to serve the merely homeostatic function of replacing neurons lost to turnover and injury ^5^. Results of the present study, together with those of a previous one ^30^, challenge these assumptions by demonstrating that neurogenesis is not entirely stochastic with respect to subtype, but rather that the birthrates of a fraction of OSN subtypes can be selectively and directionally regulated by discrete odor experiences (Figure 7). These findings suggest the possibility that persistent neurogenesis within the OE serves an unknown adaptive function in addition to the known homeostatic one. It is conceivable, for example, that the acceleration of the birthrates of specific OSN subtypes could selectively enhance sensitivity to odors detected by those subtypes by increasing their representations within the OE ^57–60^. Under this scenario, OSNs of affected subtypes might have baseline representations that lie within the dynamic range for signaling to projection neurons under physiological concentrations of cognate odors, such that accelerated neurogenesis could enhance an animal’s sensitivity to odors detected by these subtypes. This effect could have relevance to intriguing and unexplained observations in both rodents and humans that exposure to specific odors can dramatically increase sensitivity to them ^61–66^. Alternatively, or in addition, OSNs produced *via* odor-dependent neurogenesis could conceivably enable the formation of new OB glomeruli and synaptic connections with projection neurons ^67–70^. Under this scenario, stimulation-dependent neurogenesis of specific subtypes could alter inputs to the olfactory cortex and thereby regulate the perception of, and behavioral responses to, specific odors.

**Figure 7.**
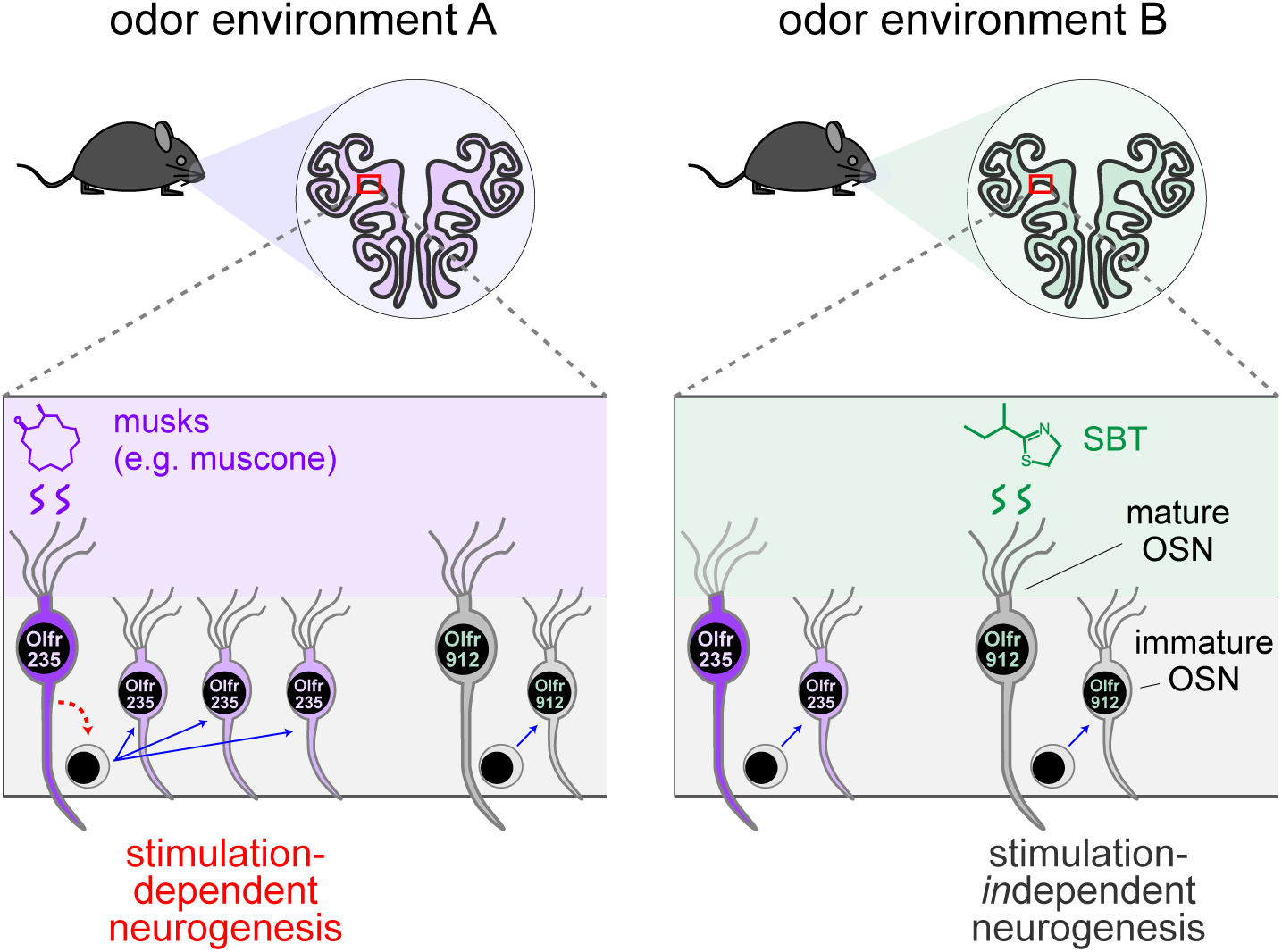
Model for how specific odors selectively increase quantities of newborn OSNs of subtypes that are stimulated by them. A fraction of OSN subtypes (e.g., Olfr235), upon stimulation by discrete odors (e.g., musks), undergo accelerated rates of neurogenesis. Most subtypes (e.g., Olfr912) do not exhibit altered rates of neurogenesis upon stimulation by odors that stimulate them (e.g., SBT). One hypothetical mechanism involves selective signaling from odor-stimulated mature OSNs of specific subtypes to neural progenitors.

### A special role for musk-responsive OSN subtypes

Results of the present study demonstrate that the birthrates of musk-responsive subtypes can be regulated by exposure to musk odors, a group of molecules that are naturally emitted by numerous mammalian species ^71–73^. For some mammals, musk odors are known to function in attracting mates, marking territory, and deterring predators ^55,74^. Moreover, exposure to musk odors has been reported to cause physiological changes in some mammals, including humans, suggesting that they can function as semiochemicals ^75,76^. In mice, the physiological functions of musk odors, if any, are unknown, although they have been found to be selectively attractive to male mice ^77^. Previous findings that exogenous musk odors activate a small number of related and evolutionarily conserved mouse ORs ^32–34^ and that odors emitted selectively by male mice activate OSN subtypes that express a subset of these ORs ^19^ suggest the possibility that mice also emit musk-like molecules. Our preliminary findings that mouse preputial gland extracts contain molecules that are structurally related to known musk odors provide additional support for this possibility.

Our findings that the exposure of mice to odors from adolescent males, but not females, accelerates the birthrate of Olfr235 OSNs, provide a mechanistic explanation for previous observations that mice exposed to male odors exhibit higher representations of this subtype ^19,24^. Curiously, two other musk-responsive subtypes that displayed a higher representation in males and females housed with males, Olfr1440 and Olfr1431 ^19^, showed stimulation-dependent changes in newborn OSN quantities in both adolescent male and female mice, while a third such subtype, Olfr1437, exhibited no occlusion-mediated changes in quantities of newborn OSNs in adolescent mice of either sex (Figures 1, 2, and data not shown). Considering the close relationship of the ORs that define musk-responsive OSN subtypes ^32^, these differences are intriguing. One conceivable explanation is that musk-responsive OSN subtypes vary in their sensitivity to distinct musk-like odorant molecules ^33,77^, which may be differentially emitted by mice in an age- and sex-dependent manner. Indeed, mouse odor profiles are known to vary considerably as a function of age and sex ^47–50^. In support of this, we have found that the exposure of adolescent mice to adults selectively intensifies UNO-induced open-side biases in quantities of newborn Olfr1431 OSNs. A more complete understanding of the role of odor-stimulation-dependent neurogenesis in altering the representations of distinct musk-responsive subtypes in the mouse OE will require determining the identities and sources of the natural ligands that stimulate these subtypes and how exposure to these odorants affects OSN birthrates. Future studies will also be needed to determine the mechanism that endows musk-responsive subtypes, as well as a fraction of other subtypes whose ligands have yet to be identified ^30^, with the capacity to undergo stimulation-dependent neurogenesis, and whether these subtypes detect odors with special functional salience.

## Acknowledgements

We are grateful to members of the Santoro lab for helpful discussions and for comments and suggestions on this manuscript. This article is based upon work supported by the National Science Foundation (Grant No. 1943528) and the National Institutes of Health (NIDCD; R01DC019936).

## Author contributions

Conceptualization, S.W.S., K.H.; Methodology, S.W.S., K.H.; Investigation, K.H., M.S., K.R., S.W.S.; Formal analysis, K.H., S.W.S., M.S.; Data curation, R.O., K.H., S.W.S.; Writing, S.W.S., K.H.; Funding acquisition, S.W.S.; Supervision, S.W.S.

---

**Appendix 1–figure 1.**
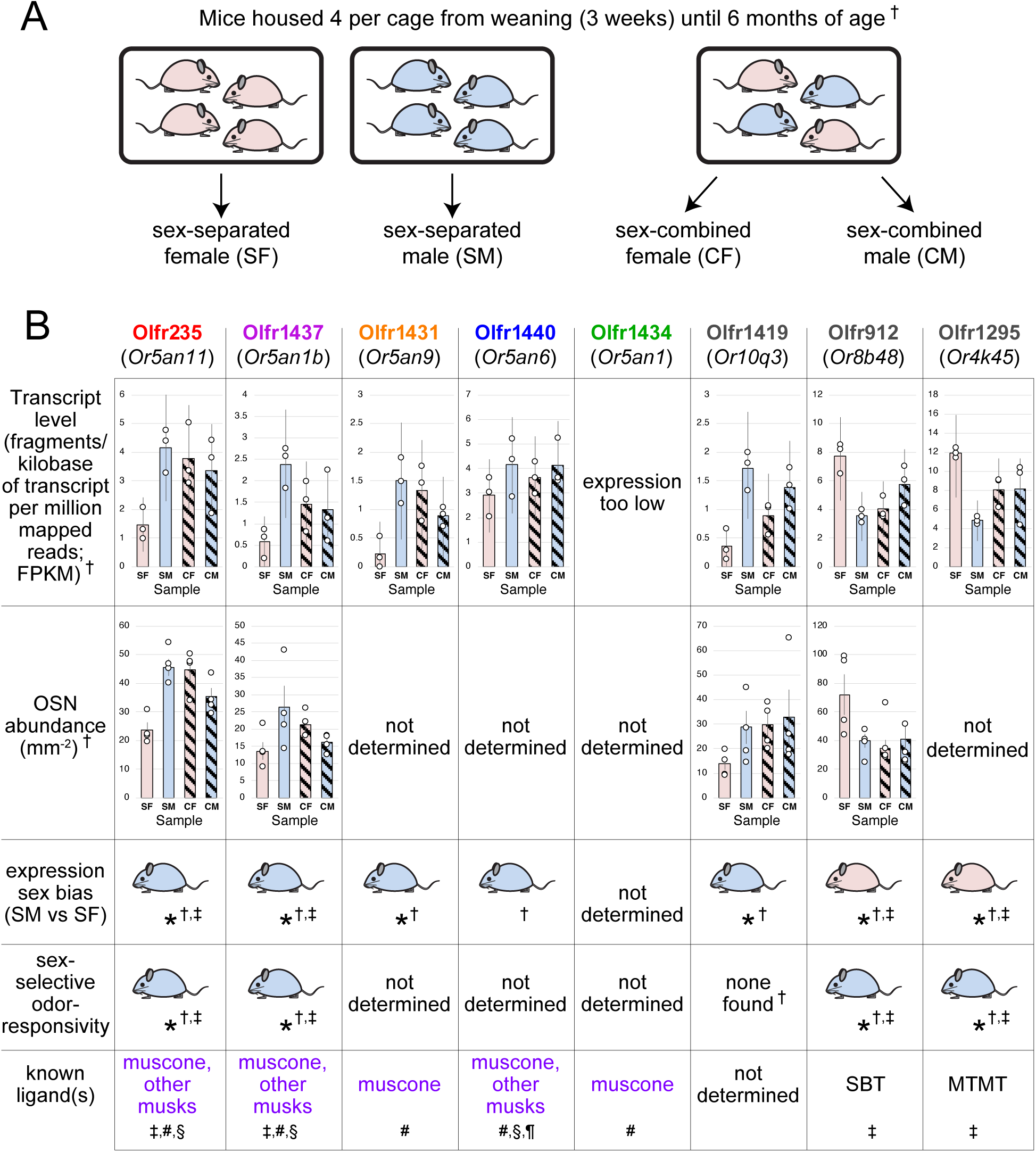
Identification of OSN subtypes that are candidates for undergoing sex-specific- and/or musk odor-accelerated neurogenesis. A. In a previous study, the OE transcript profiles of female and male mice that were housed either sex-separated or sex-combined from weaning (PD 21) until 6 months of age were profiled and compared *via* bulk RNA-seq ^19,78^. B. OSN subtypes previously identified as responsive to sex-specific odors and/or musk-like odors. SBT, 2-sec-butyl-4,5-dihydrothiazole; MTMT, (methylthio)methanethiol; †, ^19,78^; ‡, ^24^; #, ^32^; §, ^33^; ¶, ^34^. Error bars: 95% confidence intervals.

**Figure 1–figure supplement 1.**
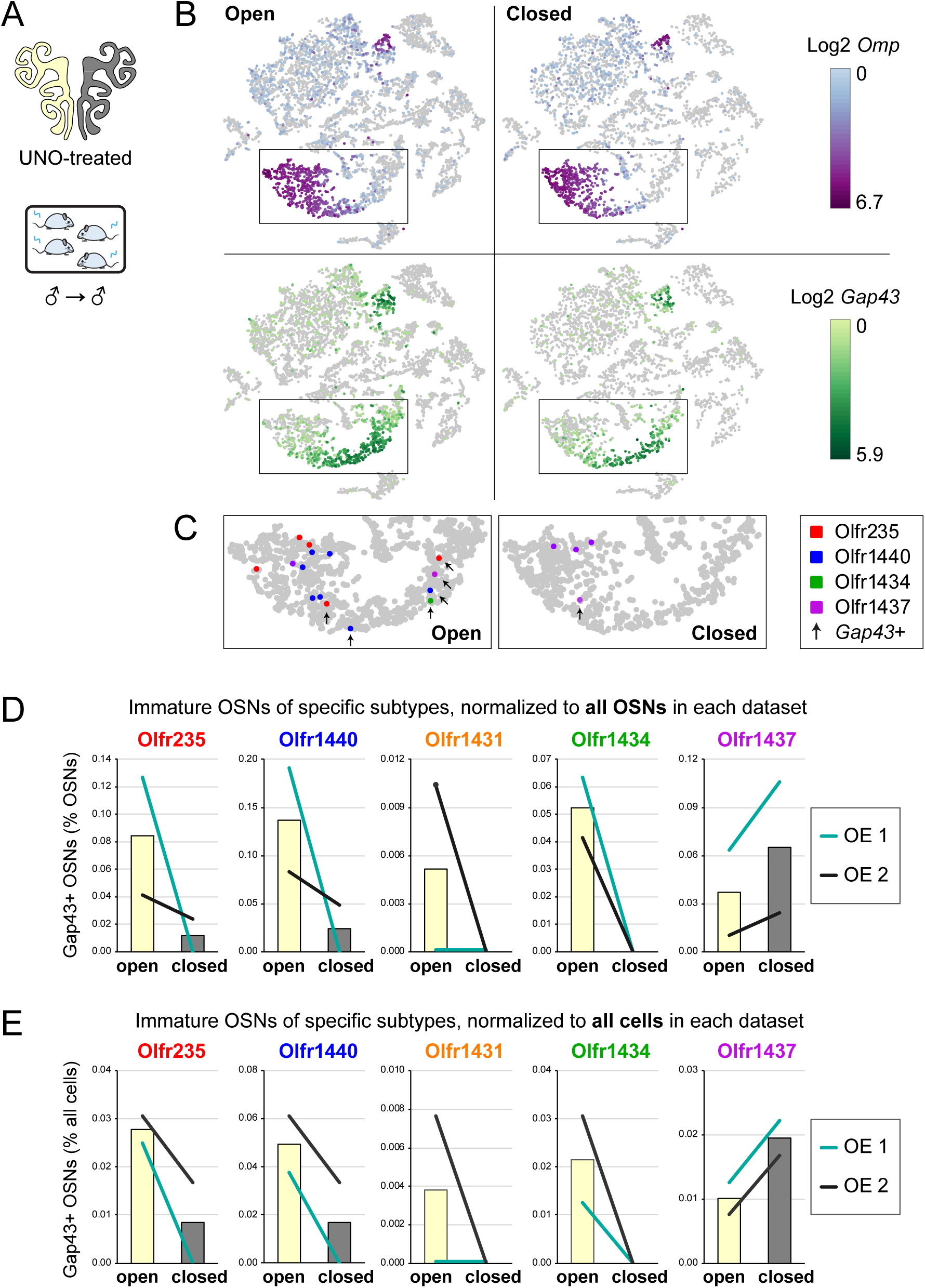
scRNA-seq analyses of the open and closed sides of whole OEs from UNO-treated adolescent male mice reveal greater quantities of newborn OSNs of musk-responsive subtypes on the open side of the OE relative to the closed. A. Experimental conditions used to generate scRNA-seq datasets for assessing the effects of UNO on quantities of newborn OSNs of specific subtypes in the OEs of adolescent male mice. B. *t*-distributed stochastic neighbor embedding (*t*-SNE) representations of all cells within scRNA-seq datasets corresponding to the open (*left*) and closed (*right*) sides of a male mouse OE (OE 1; generated in a previous study ^30^), with *Omp* (mOSNs; *top*) and *Gap43* (iOSNS; *bottom*) expression shown. OE 1 datasets were generated as outlined in Figure 1 (UNO-treated at PD 14, weaned sex-separated at P21, and sacrificed at PD 28). C. Enlarged view of the region of the *t*-SNE plot outlined in panel B, with OSNs of known musk-responsive subtypes indicated by colored dots, and immature (*Gap43*+) OSNs indicated by black arrows. D, E. Quantifications of immature (*Gap43*+) OSNs of musk-responsive subtypes as percentages of all OSNs (D) or all cells (E) within the OE 1 and OE 2 (generated in this study) datasets (*lines*), with mean values for the two datasets indicated (*bars*). In the OE 1 dataset, OSNs represent 943 of 4501 (21%) and 1573 of 7990 (20%) of all cells on the closed and open sides, respectively. In the OE 2 dataset, OSNs represent 8271 of 11919 (66%) and 9649 of 13101 (71%) of all cells on the closed and open sides, respectively.

**Figure 1–figure supplement 2.**
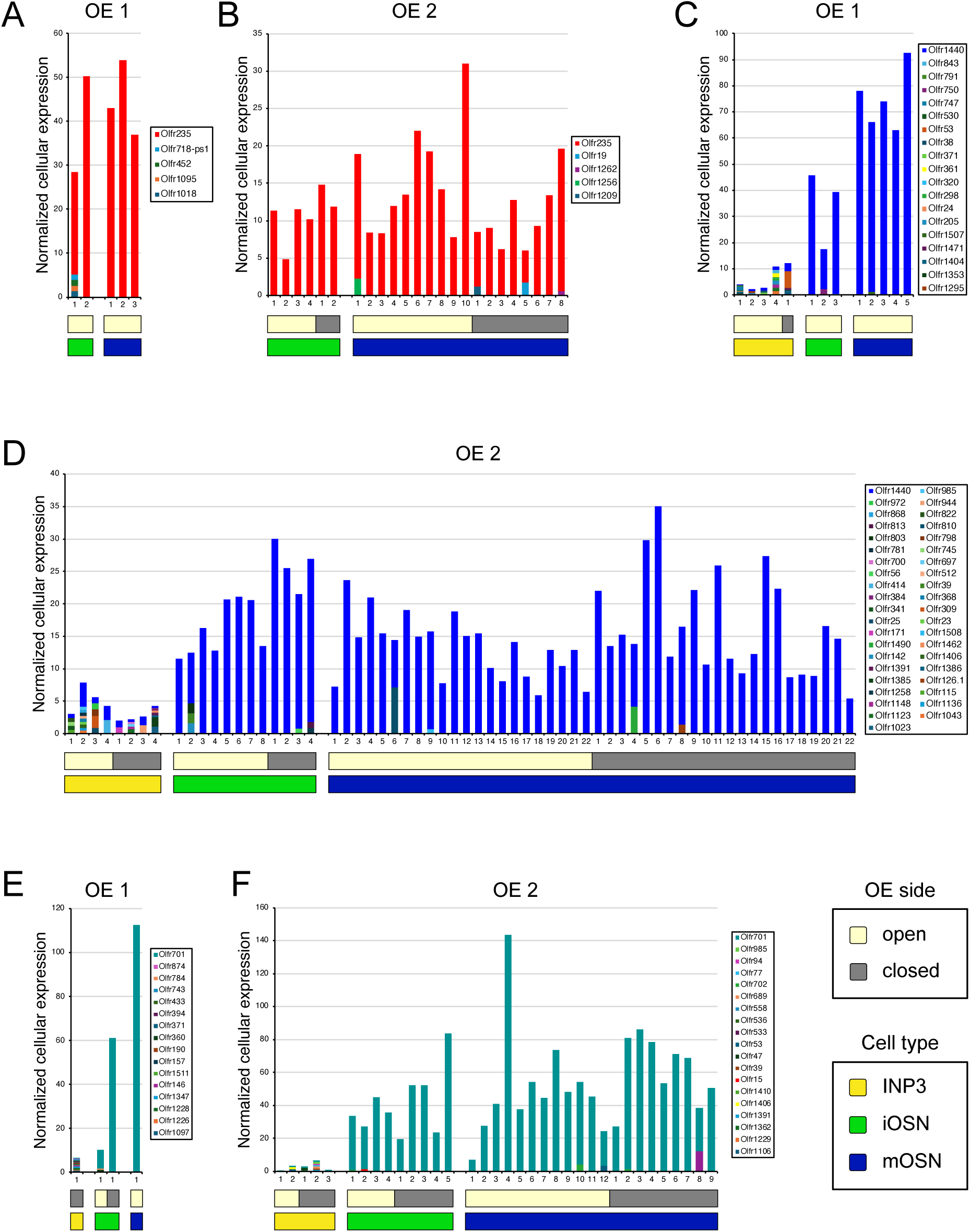
Individual OSNs of musk-responsive subtypes on the open and closed sides of OEs from UNO-treated male mice exhibit typical OR expression within cells of the OSN lineage. A-F. Cellular OR transcript levels (normalized to all transcripts within each cell) within INP3 (*horizontal dark yellow bars*), iOSN (*horizontal bright green bars*), and mOSN (*horizontal dark blue bars*) cells that contain transcripts encoding the musk responsive ORs Olfr235 (A, B) and Olfr1440 (C, D), and the randomly selected OR Olfr701 (*Or2ag2b*) from scRNA-seq datasets corresponding to 2 different male mice (OE 1, generated in a previous study ^30^, and OE 2, this study). As observed in previous studies ^38,40–45^, INP3 cells exhibit low levels of polygenic OR transcripts, while iOSNs and mOSNs exhibit high levels of monogenic OR transcripts. Moreover, OSN lineage cells of the same subtype and stage of maturity exhibit similar OR transcript levels and monogenicity within the open (*light yellow bars*) and closed sides (*gray bars*) of the OEs. Note: Olfr235 transcripts were not identified within INP3 cells of either dataset.

**Figure 2–figure supplement 1.**
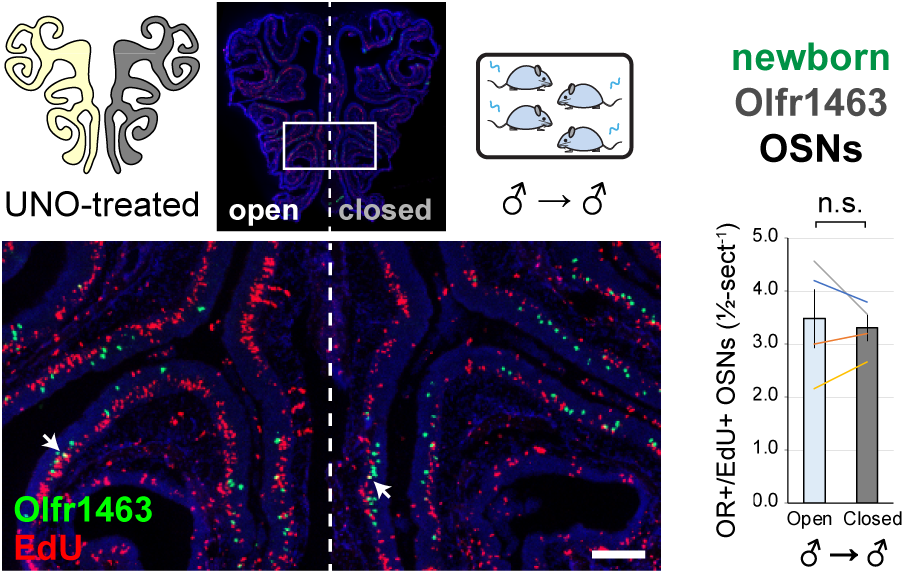
EdU birthdating analyses show that UNO-treated adolescent male mice do not exhibit significantly different quantities of newborn OSNs of control subtype Olfr1463 on the open side of the OE relative to the closed. OE tissue samples used for assessing the effects of UNO on quantities of newborn OSNs of subtype Olfr1463 were generated as outlined in Figure 2. OEs were sectioned and analyzed using Olfr1463-specific RNA-FISH and EdU staining. *Left*: Representative image of OE sections from a UNO-treated adolescent male mouse that was exposed, at the time of EdU labeling, to male littermates (♂ → ♂), with newborn OSNs (OR+/EdU+) indicated by white arrows. *Right*: Quantifications of newborn OSNs on the open and closed sides of tissue sections spanning the anterior-posterior lengths of OEs from UNO-treated male mice reveal no significant open-side bias in quantities of newborn OSNs of subtype Olfr1463. Scale bar: 150 µm. Each line represents a distinct mouse (*n* = 4 mice [≥ 5 sections/mouse]). n.s. *P >* 0.05; ratio paired two-tailed t-test. Error bars: SEM.

**Figure 2–figure supplement 2.**
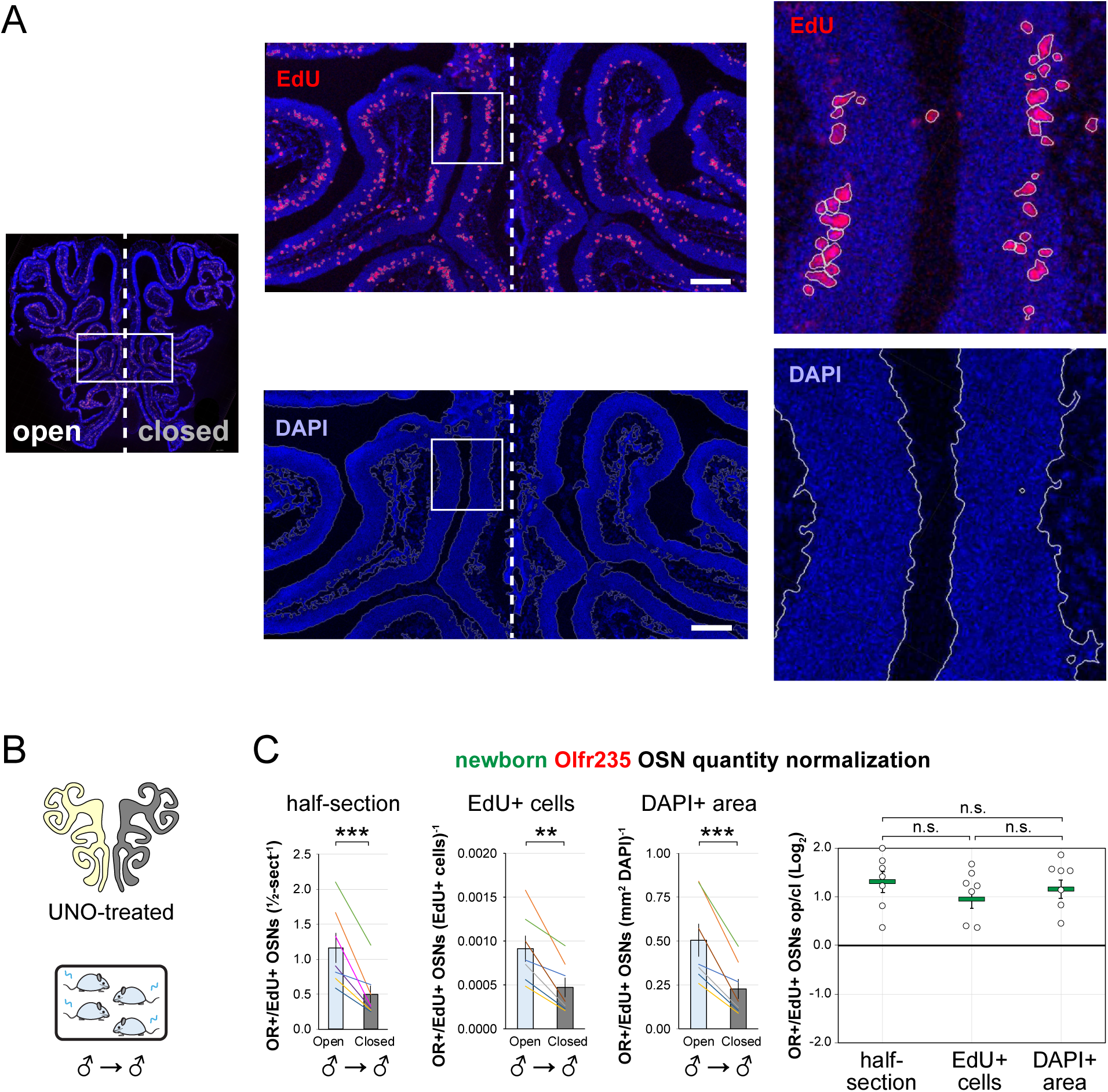
Comparison of normalization methods for assessing the effects of exposure to male odors on quantities of newborn Olfr235 OSNs in the OEs of UNO-treated male mice. A. Representative image corresponding to an EdU- and DAPI-stained OE section from an adolescent male mouse that was generated as outlined in Figure 2. EdU+ nuclei (*top*) and DAPI+ regions (*bottom*) are outlined in white, with medium (*middle*), and high magnification (*right*) views corresponding to the boxed regions within the preceding images. Scale bars: 150 µm. B. Experimental conditions used to generate OE tissues from UNO-treated adolescent male mice. C. Quantifications of newborn Olfr235 OSNs on the open and closed sides of tissue sections spanning the anterior-posterior lengths of OEs. OE sections were analyzed using Olfr235-specific RNA-FISH and EdU staining and newborn Olfr235 OSN quantities were normalized by half-section, number of EdU+ cells, or DAPI+ area. Each of the normalization methods yielded open-side biases of similar magnitude in quantities of newborn Olfr235 OSNs (*left*), with no significant differences in corresponding UNO effect sizes observed (*right*). Each line or circle represents a distinct mouse (*n* = 7–10 mice [≥ 5 sections/mouse] per condition). \*\*\**P* < 0.001; \*\**P* < 0.01; \**P* < 0.05; n.s. *P>* 0.05; ratio paired two-tailed t-test (C, *left*); one-way ANOVA test, FDR-adjusted (C, *right*). Error bars: SEM. Data for newborn OSN quantities normalized by half-section correspond to Figure 2.

**Figure 3–figure supplement 1.**
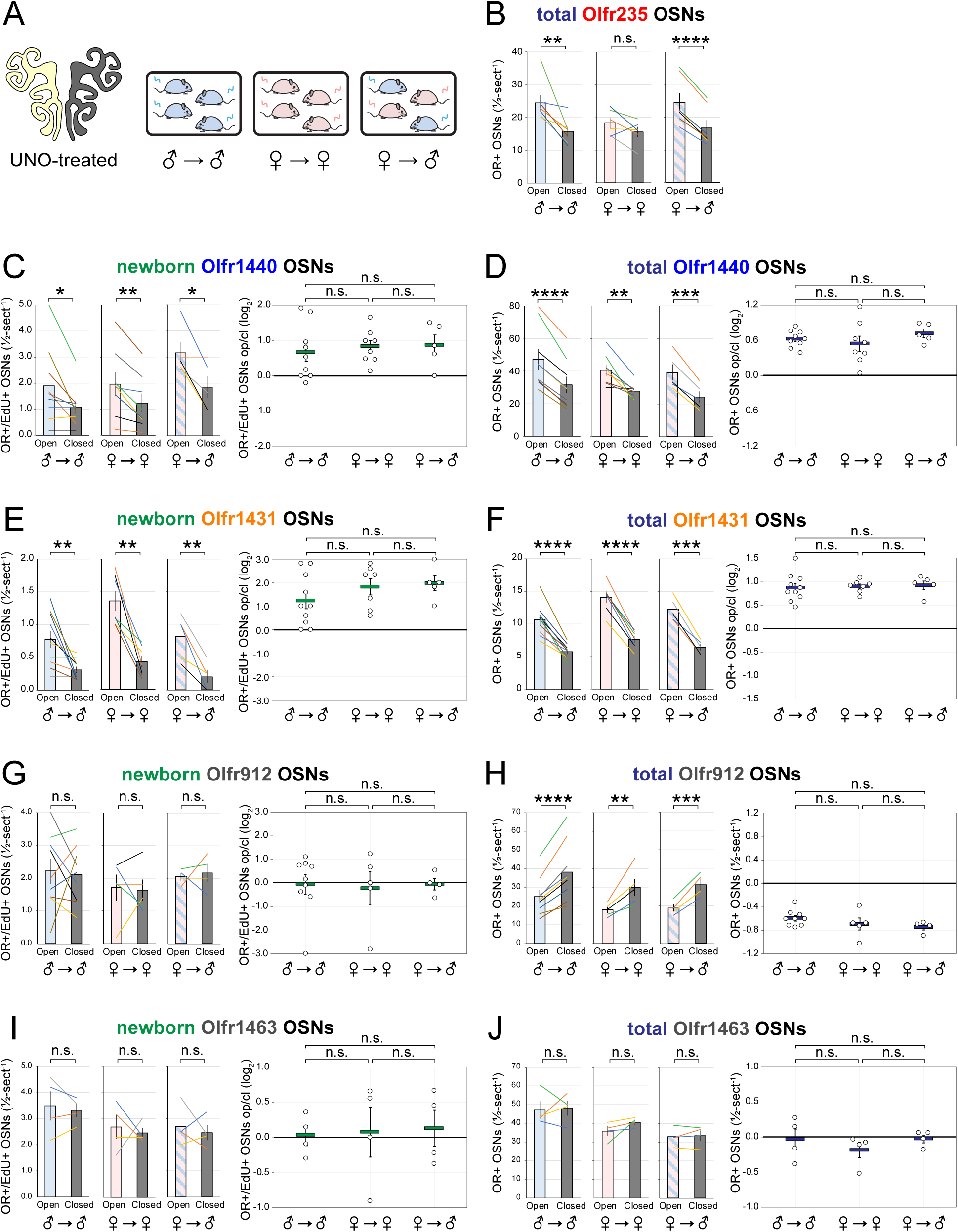
Effects of exposure to male odors on differences in quantities of newborn and total OSNs of musk-responsive and control subtypes on the open and closed sides of the OEs of UNO-treated mice. A. Experimental conditions used for tissue generation. OE tissue samples were generated according to the experimental timeline outlined in Figure 3, sectioned, and analyzed using OR-specific RNA-FISH and EdU staining. B. Quantifications of total (OR+) Olfr235 OSNs on the open and closed sides of tissue sections spanning the anterior-posterior lengths of OEs from UNO-treated mice reveal significant open-side biases in males and in females exposed to male littermates (♂ → ♂ or ♀ → ♂) but not in females exposed to just female littermates (♀ → ♀). C-F. Quantifications of Olfr1440 (C, D) and Olfr1431 (E, F) OSNs on the open and closed sides of tissue sections spanning the anterior-posterior lengths of OEs from UNO-treated mice reveal significant open-side biases (*left*) in quantities of newborn (OR+/EdU+; C, E) and total (D, F) OSNs of both subtypes under all odor conditions (♂ → ♂, ♀ → ♂, ♀ → ♀), with no significant differences in UNO effect sizes (*right*) observed. G, H. Quantifications of Olfr912 OSNs on the open and closed sides of tissue sections spanning the anterior-posterior lengths of OEs from UNO-treated mice reveal no significant open-side biases in quantities of newborn OSNs (G), but significant closed-side biases in total OSNs of this subtype (H) under all 3 conditions. Observed effects of UNO on total Olfr912 OSNs may be attributable to an increased rate of cell death for mature OSNs of this subtype due to overstimulation in the presence of male odors ^19,24^, to which all mice were exposed during at least part of the UNO period. I, J. Quantifications of Olfr1463 OSNs on the open and closed sides of tissue sections spanning the anterior-posterior lengths of OEs from UNO-treated mice reveal no significant open-side biases in quantities of newborn (I) or total (J) OSNs of this subtype under any of the conditions. Each line or circle represents a distinct mouse (*n* = 4–10 mice [≥ 5 sections/mouse] per OSN subtype and condition). \*\*\*\**P* < 0.0001; \*\*\**P* < 0.001; \*\**P* < 0.01; \**P* < 0.05; n.s. *P >* 0.05; ratio paired two-tailed t-test (B, C-J, *left*); one-way ANOVA test, FDR-adjusted (C-J, *right*). Error bars: SEM. Data for newborn OSN quantities in ♂ → ♂ samples (C, E, G, I) correspond to Figure 2 and Figure 2-figure supplement 1.

**Figure 3–figure supplement 2.**
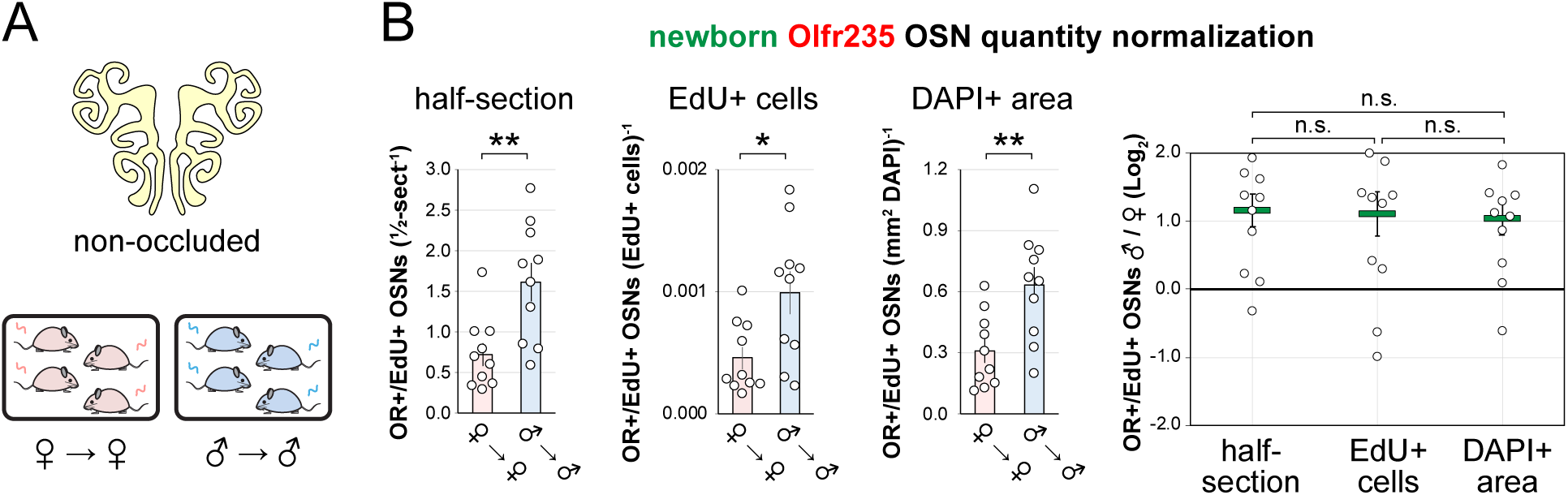
Comparison of normalization methods for assessing the effects of exposure to male odors on quantities of newborn Olfr235 OSNs in the OEs of non-occluded mice. A. Experimental conditions used to generate tissues. B. Quantifications of newborn Olfr235 OSNs within tissue sections spanning the anterior-posterior lengths of the OEs of non-occluded adolescent male and female mice that were weaned sex-separated. Newborn Olfr235 OSN quantities were normalized by half-section, number of EdU+ cells, or DAPI+ area. Each of the 3 normalization methods yielded differences of similar magnitude in quantities of newborn Olfr235 OSNs in non-occluded males exposed to male littermates (♂ → ♂) compared to females exposed to female littermates (♀ → ♀), with no significant differences observed in the corresponding odor exposure effect sizes (*right*; calculated from the ratios of normalized newborn Olfr235 OSNs within individual sex-separated male mice relative to the mean of normalized newborn Olfr235 OSNs within sex-separated females). Each line or circle represents a distinct mouse (*n* = 7–10 mice [≥ 5 sections/mouse] per condition). \*\**P* < 0.01; \**P* < 0.05; n.s. *P>* 0.05; ratio paired two-tailed t-test (B, *left*); one-way ANOVA test, FDR-adjusted (B, *right*). Error bars: SEM. Data for newborn OSN quantities normalized by half-section (B) correspond to Figure 3G.

**Figure 3–figure supplement 3.**
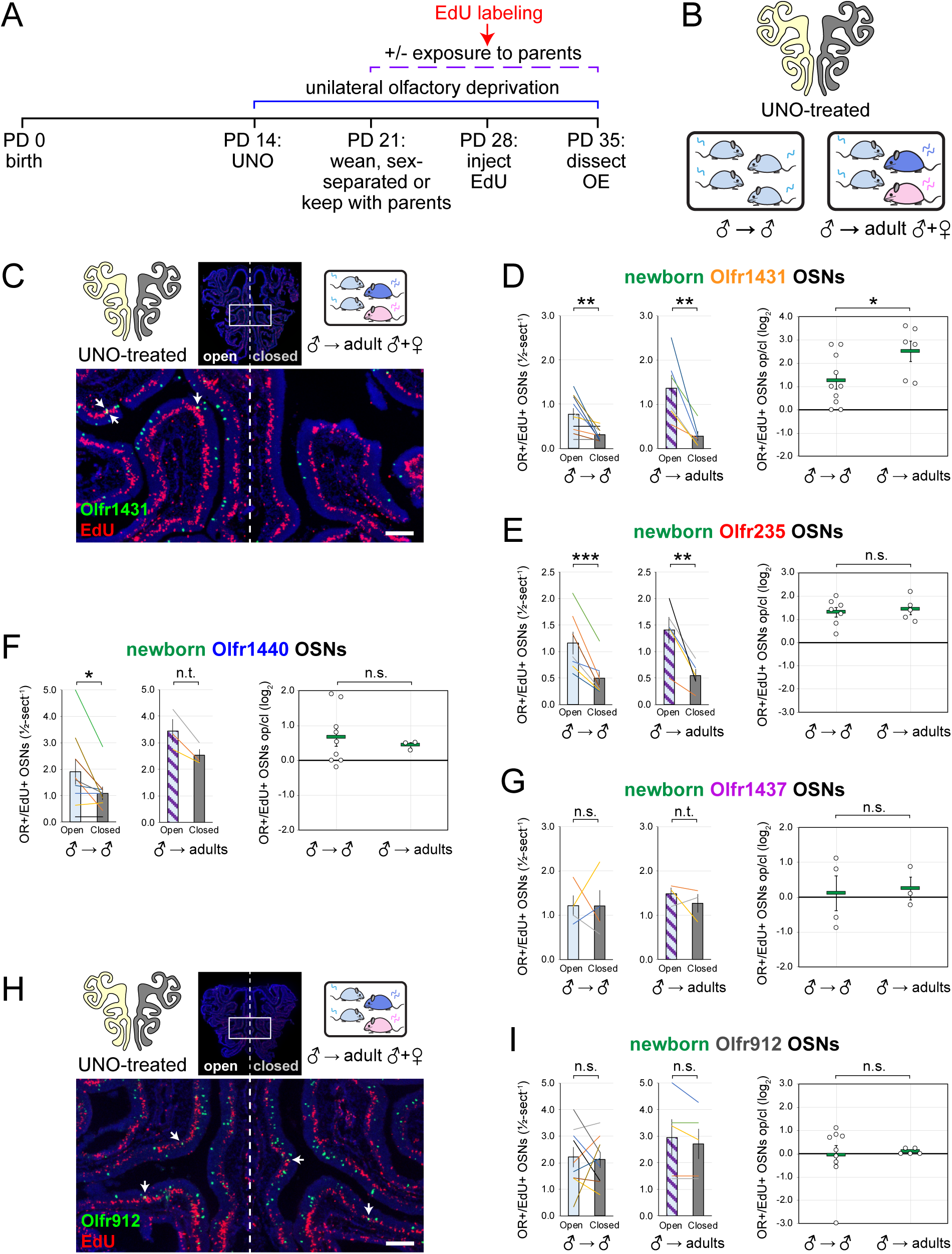
UNO-induced open-side biases in quantities of newborn Olfr1431 OSNs are increased by exposure to adult mice. A, B. Experimental timeline (A) and conditions (B) used to generate OE tissue samples for assessing the effects of exposure to adult mouse odors on quantities of newborn OSNs of specific subtypes. Male mice were UNO-treated at PD 14, weaned either sex-separated (♂ → ♂) or kept with parents (♂ → adult ♀ + ♂) at PD 21, EdU-labeled at PD 28, and sacrificed at PD 35. OEs were sectioned and analyzed using OR- specific RNA-FISH and EdU staining. C, H. Representative images of OE sections from UNO- treated adolescent male mice that were exposed, at the time of EdU labeling, to their parents, with newborn Olfr1431 (C) or Olfr912 OSNs (OR+/EdU+) indicated by white arrows. D-G, I. Quantifications of newborn Olfr1431 (D), Olfr235 (E), Olfr1440 (F), Olfr1437 (G), and Olfr912 (I) OSNs on the open and closed sides of tissue sections spanning the anterior-posterior lengths of OEs of UNO-treated mice that were exposed, at the time of EdU labeling, to just their male littermates (♂ → ♂) or also their parents (♂ → adult ♀ + ♂). Greater open-side biases (*left*) and corresponding UNO effect sizes (*right*) were observed for quantities of newborn OSNs of subtype Olfr1431 (D) in the OEs of UNO-treated male mice that were exposed to their parents compared to just littermates, while other subtypes tested were not significantly affected. Scale bars: 150 µm. Each line or circle represents a distinct mouse (*n* = 3–10 mice [≥ 5 sections/mouse] per OSN subtype and condition). \*\*\**P* < 0.001; \*\**P* < 0.01; \**P* < 0.05; n.s. *P>* 0.05; n.t. not tested (n < 4); ratio paired two-tailed t-test (D-G, I, *left*); unpaired two-tailed t-test (D-G, I, *right*). Error bars: SEM. Data for ♂ → ♂ samples correspond to Figure 2.

**Figure 4–figure supplement 1.**
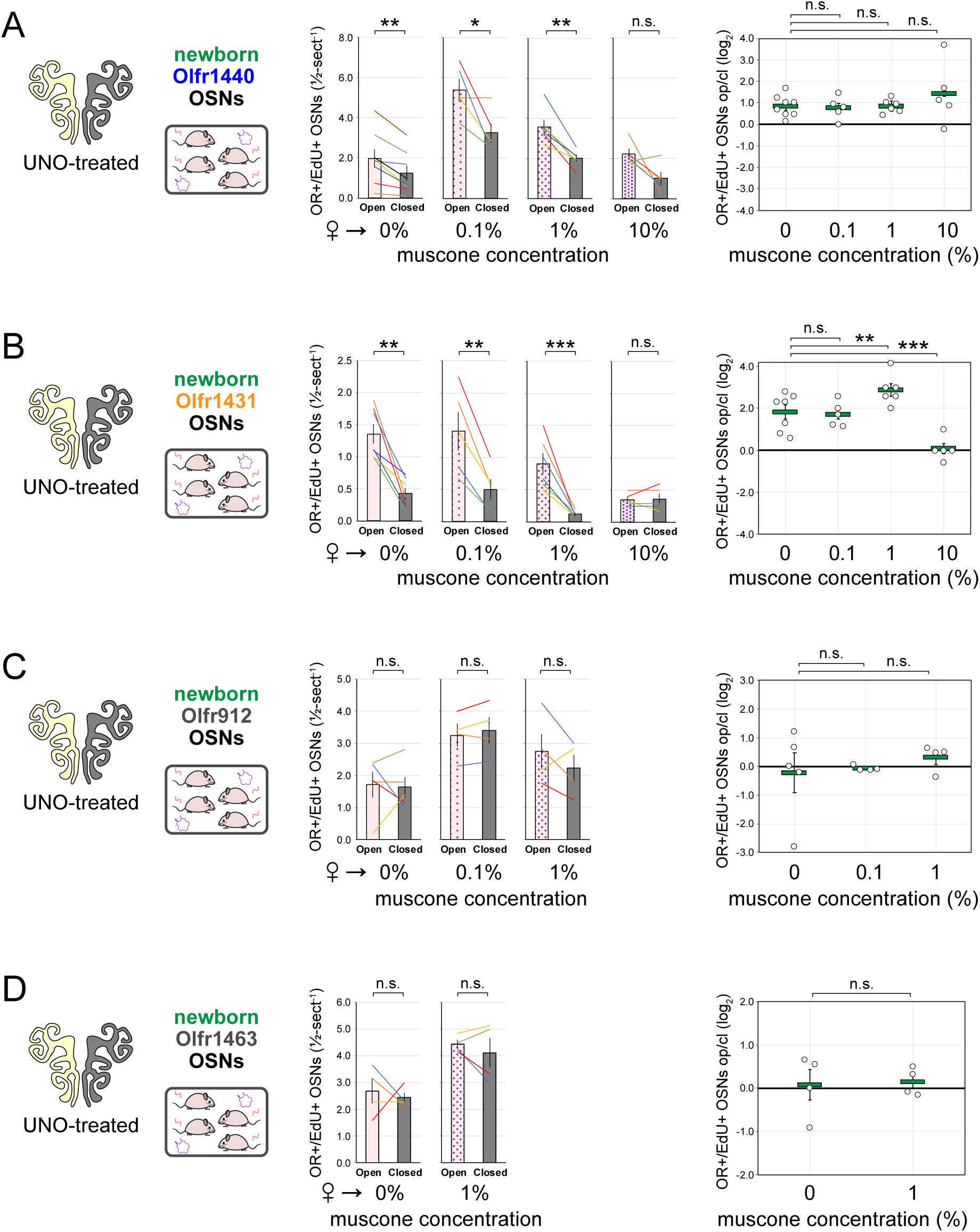
Exposure of UNO-treated female mice to muscone causes concentration-dependent differences in quantities of newborn OSNs of musk responsive subtypes on the open and closed sides of the OE. A–D. *Left*: Experimental conditions used to generate OE tissue samples for assessing the effects of exposure to muscone on quantities of newborn OSNs of specific subtypes. OE tissue samples were generated according to the experimental timeline outlined in Figure 4, sectioned, and analyzed using OR-specific RNA-FISH and EdU staining. *Middle, right*: Quantifications of newborn (OR+/EdU+) OSNs of the musk-responsive subtypes Olfr1440 (A) and Olfr1431 (B), and the control subtypes Olfr912 (C) and Olfr1463 (D) on the open and closed sides of tissue sections spanning the anterior-posterior lengths of the OEs of UNO-treated female mice that were exposed, at the time of EdU labeling, to 0, 0.1, 1, or 10% muscone. Each line or circle represents a distinct mouse (*n* = 4–8 mice [≥ 5 sections/mouse] per OSN subtype and condition). \*\*\**P* < 0.001; \*\**P* < 0.01; \**P* < 0.05; n.s. *P>* 0.05; ratio paired two-tailed t-test (*middle*); unpaired two-tailed t-test (*right*). Error bars: SEM. Data for ♀ → 0% muscone samples correspond to Figure 3–figure supplement 1.

**Figure 4–figure supplement 2.**
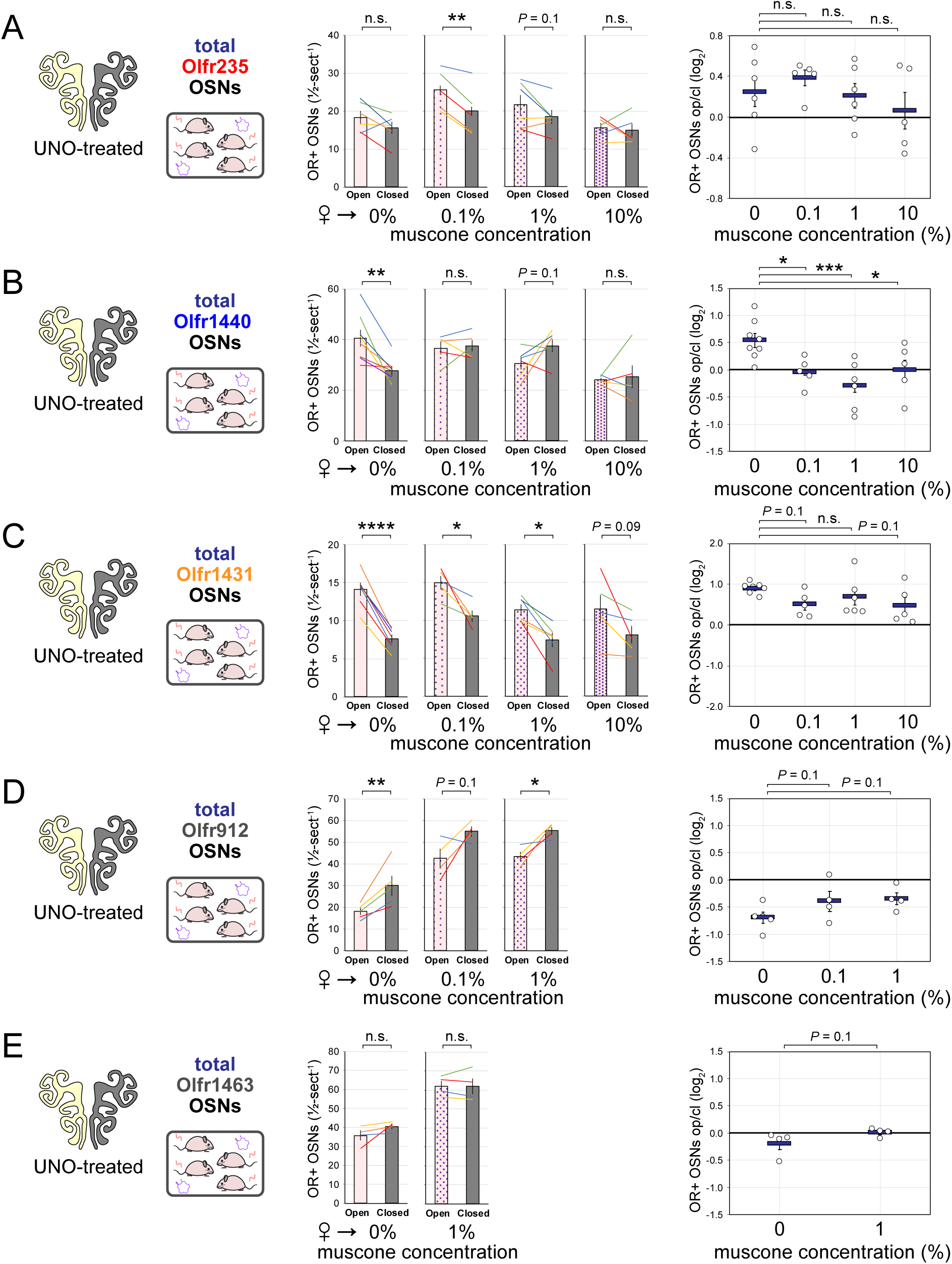
Muscone exposure induces concentration-dependent differences in quantities of total OSNs of musk responsive subtypes on the open and closed sides of the OEs of UNO-treated mice. A–E. *Left*: Experimental conditions used to generate OE tissue samples for assessing the effects of exposure to muscone on quantities of total OSNs of specific subtypes. OE tissue samples were generated according to the experimental timeline outlined in Figure 4, sectioned, and analyzed using OR-specific RNA-FISH and EdU staining. *Middle, right*: Quantifications of total (OR+) OSNs of the musk-responsive OSN subtypes Olfr235 (A), Olfr1440 (B) and Olfr1431 (C), and the control subtypes Olfr912 (D) and Olfr1463 (E) on the open and closed sides of tissue sections spanning the anterior-posterior lengths of the OEs of UNO-treated female mice that were exposed, at the time of EdU labeling, to 0, 0.1, 1, or 10% muscone. Note that the reduced quantities of total OSNs of subtype Olfr912 may be attributable to a reduced rate of survival for mature Olfr912 OSNs in the presence of male odors ^19,24^, to which mice were exposed prior to weaning. Each line or circle represents a distinct mouse (*n* = 4–8 mice [≥ 5 sections/mouse] per OSN subtype and condition). \*\**P* < 0.01; \**P* < 0.05; n.s. *P>* 0.05; two-tailed paired t-test (*middle*); two-tailed unpaired t-test (*right*). Error bars: SEM.

**Figure 4–figure supplement 3.**
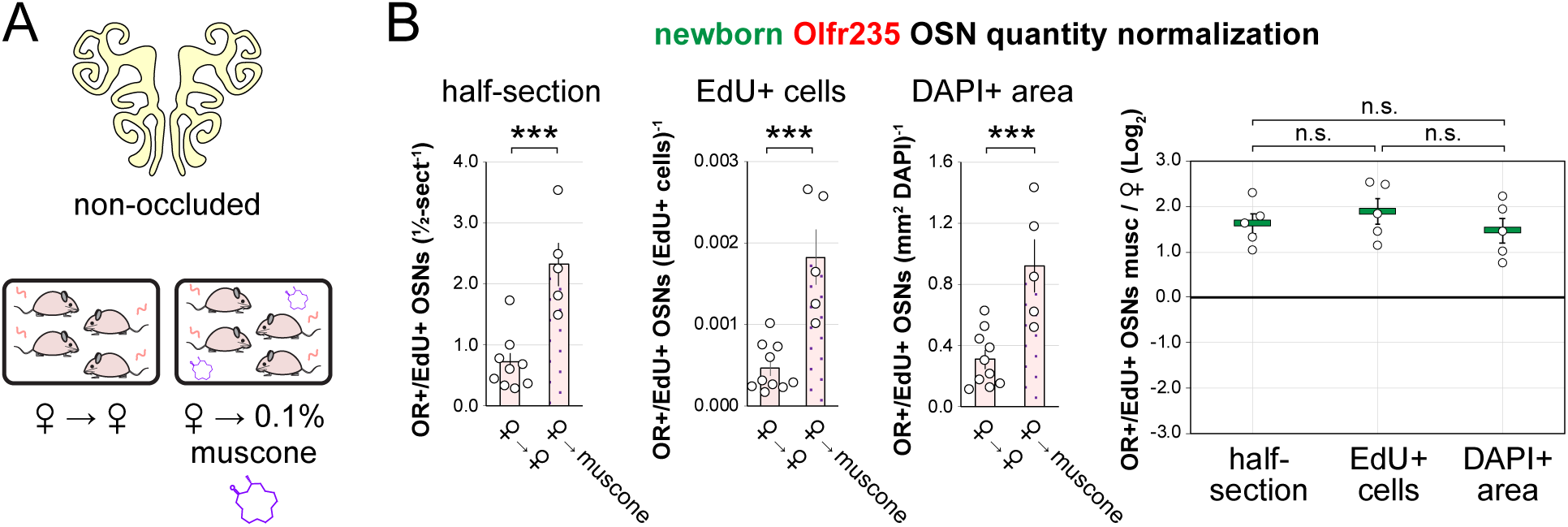
Comparison of normalization methods for assessing the effects of exposure to a musk odor on subtype-specific newborn OSN quantities in the OEs of non-occluded mice. A. Experimental conditions used to generate OE tissue samples. Samples were generated according to the experimental timeline outlined in Figure 4, sectioned, and analyzed using OR-specific RNA-FISH and EdU staining. B. Quantifications of newborn Olfr235 OSNs (OR+/EdU+) within tissue sections spanning the anterior-posterior lengths of the OEs of non-occluded female mice that were either exposed to 0.1% muscone (♀ → muscone) or unexposed to an exogenous odorant (♀ → ♀) starting from PD 21, EdU-labeled at PD 28, and sacrificed at PD 35. OE sections were analyzed using Olfr235-specific RNA-FISH and EdU staining and newborn Olfr235 OSN quantities were normalized by half-section, number of EdU+ cells, or DAPI+ area. Each of the normalization methods yielded significant differences in quantities of newborn Olfr235 OSNs in muscone-exposed compared to unexposed mice (B-*left*), with no significant differences observed in the corresponding odor exposure effect sizes (B-*right*; calculated from the ratios of normalized newborn Olfr235 OSNs within individual odor-exposed mice relative to the mean of normalized newborn Olfr235 OSNs within control mice). Each circle represents a distinct mouse (*n* = 5–10 mice [≥ 5 OE sections/mouse] per condition). \*\*\**P* < 0.001; n.s. *P>* 0.05; unpaired two-tailed t-test (B-*left*); one-way ANOVA test, FDR-adjusted (B-*right*). Error bars: SEM. Data for newborn OSN quantities normalized by half-section correspond to Figure 4D.

**Figure 4–figure supplement 4.**
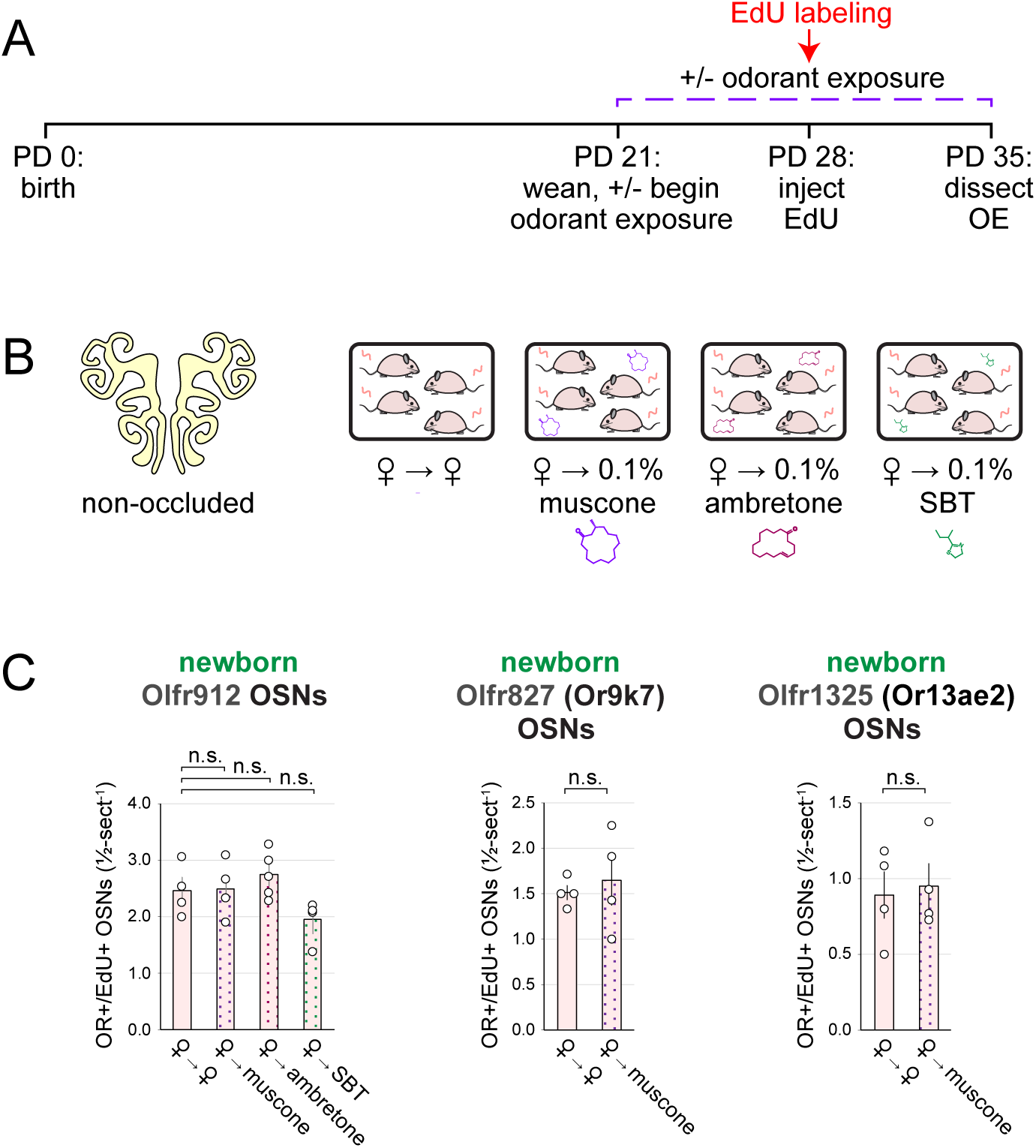
Effects of exposure of mice to musk and non-musk odors on quantities of newborn OSNs of non-musk-responsive subtypes, including those previously found to undergo stimulation-dependent changes in OSN birthrate. A, B. Experimental timeline (A) and conditions (B) used to generate OE tissue samples for assessing the effects of exposure to musk (muscone, ambretone) or non-musk (SBT) odors on quantities of newborn OSNs of specific subtypes. Non-occluded female mice were weaned sex-separated and either exposed or unexposed to an exogenous musk or non-musk odorant starting at PD 21, EdU-labeled at PD 28, and sacrificed at PD 35. OEs were sectioned and analyzed using OR-specific RNA-FISH and EdU staining. C. *Left*: Quantifications of newborn OSNs of the SBT-responsive subtype Olfr912 within tissue sections spanning the anterior-posterior lengths of OEs reveal no significant differences in non-occluded females that were exposed, at the time of EdU labeling, to a musk odorant (♀ → muscone; ♀ → ambretone) or to SBT (♀ → SBT) compared to those exposed to just female littermates (♀ → ♀). *Middle, right*: Quantifications of newborn OSNs of two subtypes previously found to undergo stimulation-dependent changes in birthrates, Olfr827 (*middle*) and Olfr1325 (*right*) ^30^, reveal no significant differences in non-occluded females that were exposed to a musk odorant compared to those exposed to just female littermates. Each circle represents a distinct mouse (*n* = 3–4 mice [≥ 5 OE sections/mouse] per OSN subtype and condition). n.s. *P>* 0.05; one-way ANOVA test, FDR-adjusted (C-*left*); unpaired two-tailed t-test (C-*middle, right*). Error bars: SEM. Data for newborn Olfr912 OSNs in ♀ → ♀ samples correspond to Figure 3H.

**Figure 6–figure supplement 1.**
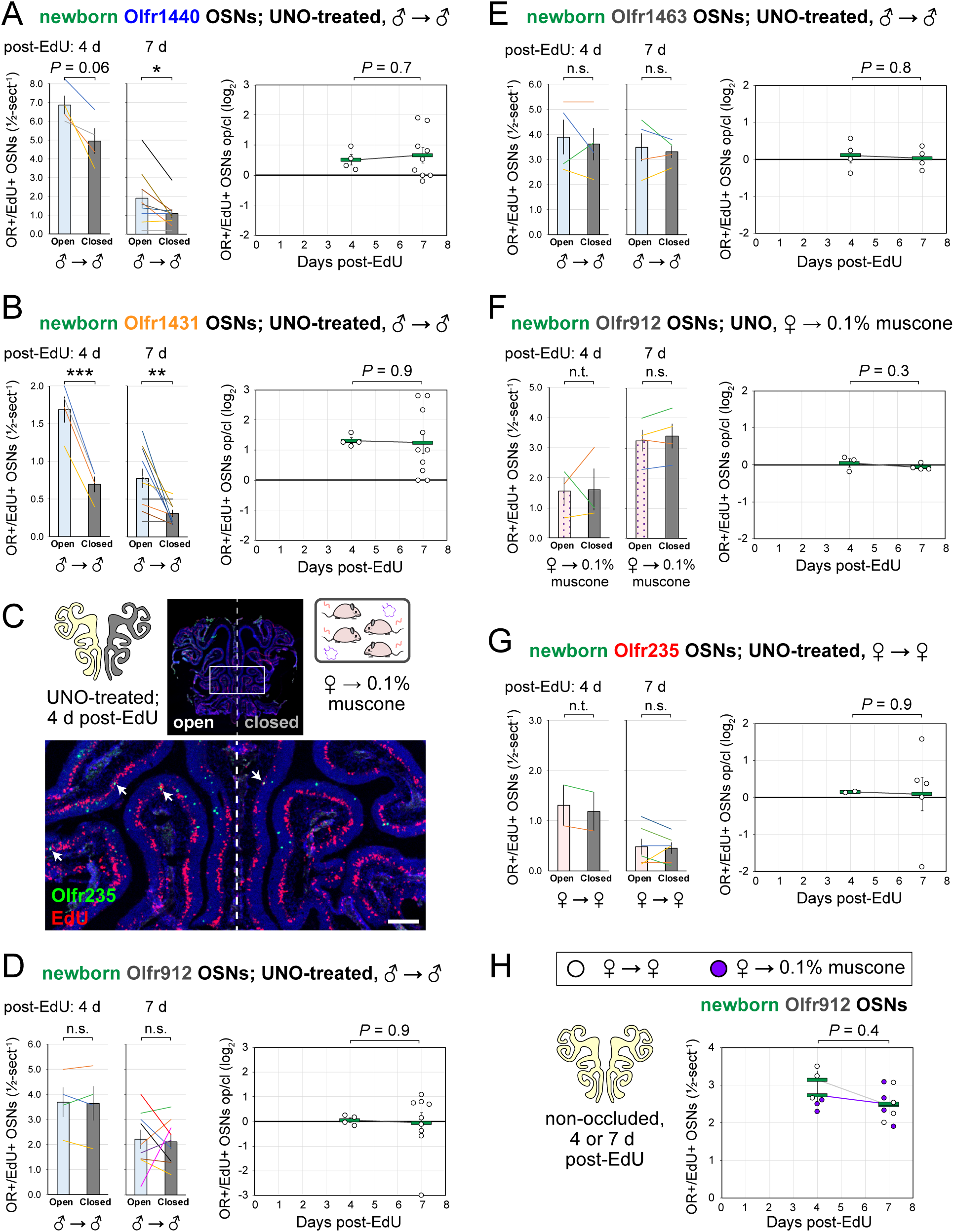
Stimulation-dependent increases in quantities of newborn OSNs of musk-responsive subtypes are stable over time following neurogenesis, consistent with a mechanism involving altered birthrate. OE tissue samples were generated according to the experimental timeline and conditions outlined in Figure 6, sectioned, and analyzed using OR- specific RNA-FISH and EdU staining. A, B, D–G. Quantifications of newborn Olfr1440 (A), Olfr1431 (B), Olfr912 (D, F), Olfr1463 (E), and Olfr235 (G) OSNs (OR+/EdU+) on the open and closed sides of tissue sections spanning the anterior-posterior lengths of OEs of UNO-treated males exposed to male littermates (♂ → ♂) (A, B, D, E), females exposed to muscone (♀ → 0.1% muscone) (F), or females exposed to female littermates (♀ → ♀) (G). Under all conditions, similar open-side biases in quantities of newborn OSNs were observed at both 4 and 7 d post-EdU, with no significant differences in UNO effect sizes observed over time. C. Representative image of OE sections from a UNO-treated female mouse that was exposed to muscone (♀ → 0.1% muscone) and sacrificed 4 d post-EdU, with newborn Olfr235 OSNs (OR+/EdU+) indicated by white arrows. Greater numbers of newborn Olfr235 OSNs were observed on the open side compared to the closed side at this timepoint. Scale bar: 150 µm. H. Quantifications of newborn OSNs of the SBT-responsive subtype Olfr912 within OEs of non-occluded female mice that were either exposed just to female littermates (♀ → ♀) or also to muscone (♀ → 0.1% muscone) and sacrificed 4 or 7 d post-EdU. No significant changes in newborn OSN quantity differences were observed between 4 and 7 d post-EdU. Each line or circle represents a distinct mouse (*n* = 2–10 mice [≥ 5 OE sections/mouse] per OSN subtype and condition). \*\*\**P* < 0.001; \*\**P* < 0.01; \**P* < 0.05; n.s. *P* > 0.05; n.t. not tested (n < 4); ratio paired two-tailed t-test (A, B, D–G-*left*); unpaired two-tailed t-test (A, B, D–G-*right*); two sample ANOVA - fixed-test, using F distribution (right-tailed; H). Error bars: SEM. Data for 7 d post-EdU samples correspond to Figures 2–4.

**Appendix 2–figure 1.**
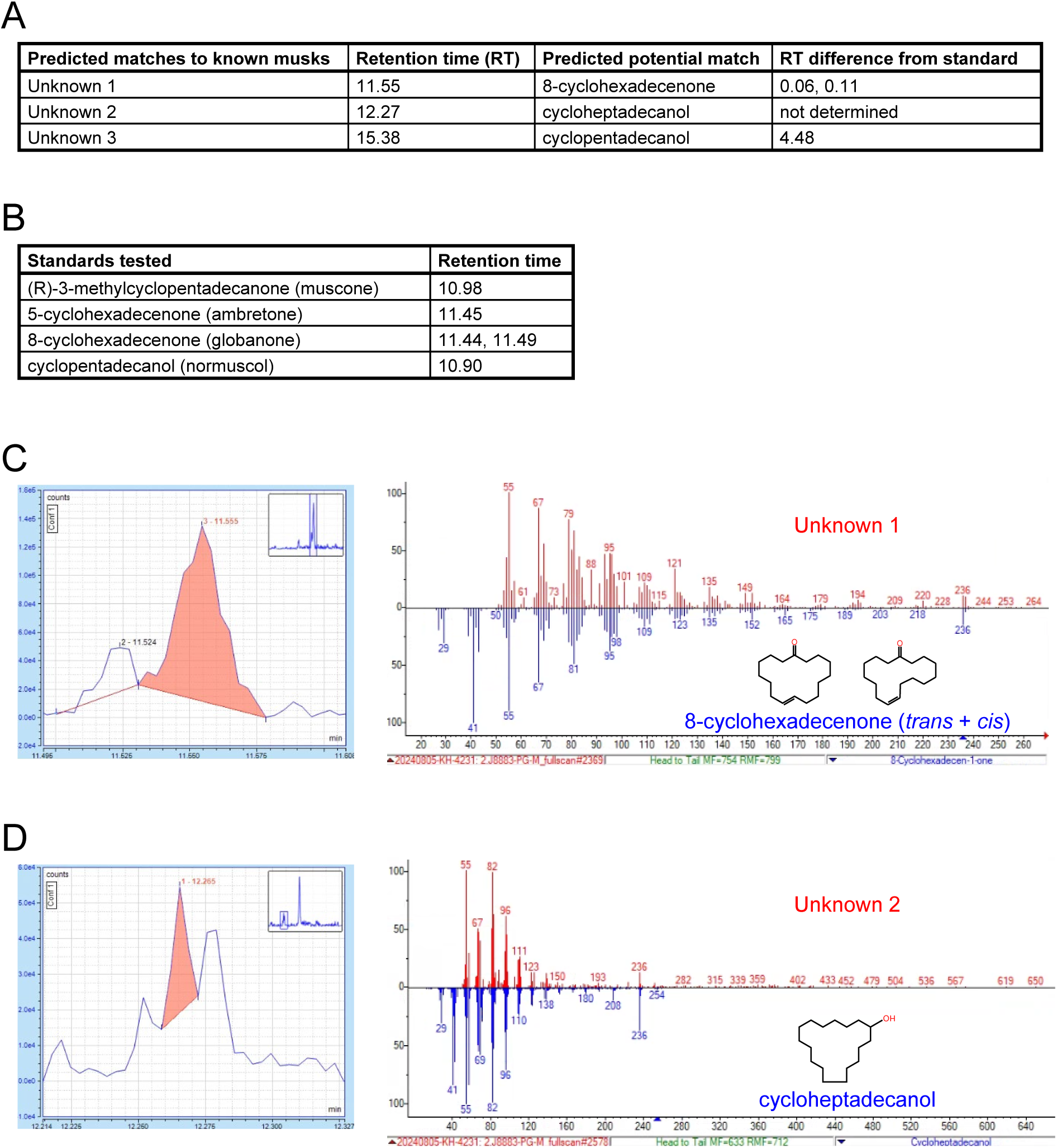
Gas chromatography – mass spectrometry (GC-MS) analyses of mouse preputial gland extracts for molecules from with structural similarity to known musk odorants. A, B. GC-MS signals from male mouse preputial gland extracts (A) and commercially available samples of 4 known musk compounds (B). Analyses of preputial glands revealed signals (Unknowns 1-3) with predicted potential matches to the indicated odorants based on spectral similarities (A). Experimental differences in retention times between unknowns and standards are indicated. C. *Left*: Region of the extracted ion chromatograph (m/z 236) of a preputial gland extract, with the signal corresponding to Unknown 1 highlighted in red. *Right*: Mass spectra corresponding to Unknown 1 (*red*) and a predicted match, 8-cyclohexadecenone (*blue*), a musk compound that was previously found to activate Olfr235 and Olfr1440 ^33^. The retention times of Unknown 1 and 8-cyclohexadecenone differ by 0.06 and 0.11 minutes (possibly corresponding to the *cis* and *trans* isomers) (A), indicating potential structural similarity. D. *Left*: Region of the extracted ion chromatograph (m/z 236) of a preputial gland extract, with the signal corresponding to Unknown 2 highlighted in red. *Right*: Mass spectra corresponding to Unknown 2 (*red*) and a predicted match, cycloheptadecanol (*blue*). Confirmation of a match will require comparison of the observed retention time of Unknown 2 (12.27 minutes) with that of a cycloheptadecanol standard (not determined). Although Unknown 3 exhibited a predicted match to cyclopentadecanol, a musk compound previously found to activate Olfr235 and Olfr1440 ^33^, the observed retention time difference of 4.48 minutes (A) indicates substantial structural dissimilarity.

**Appendix 3–table 1.**
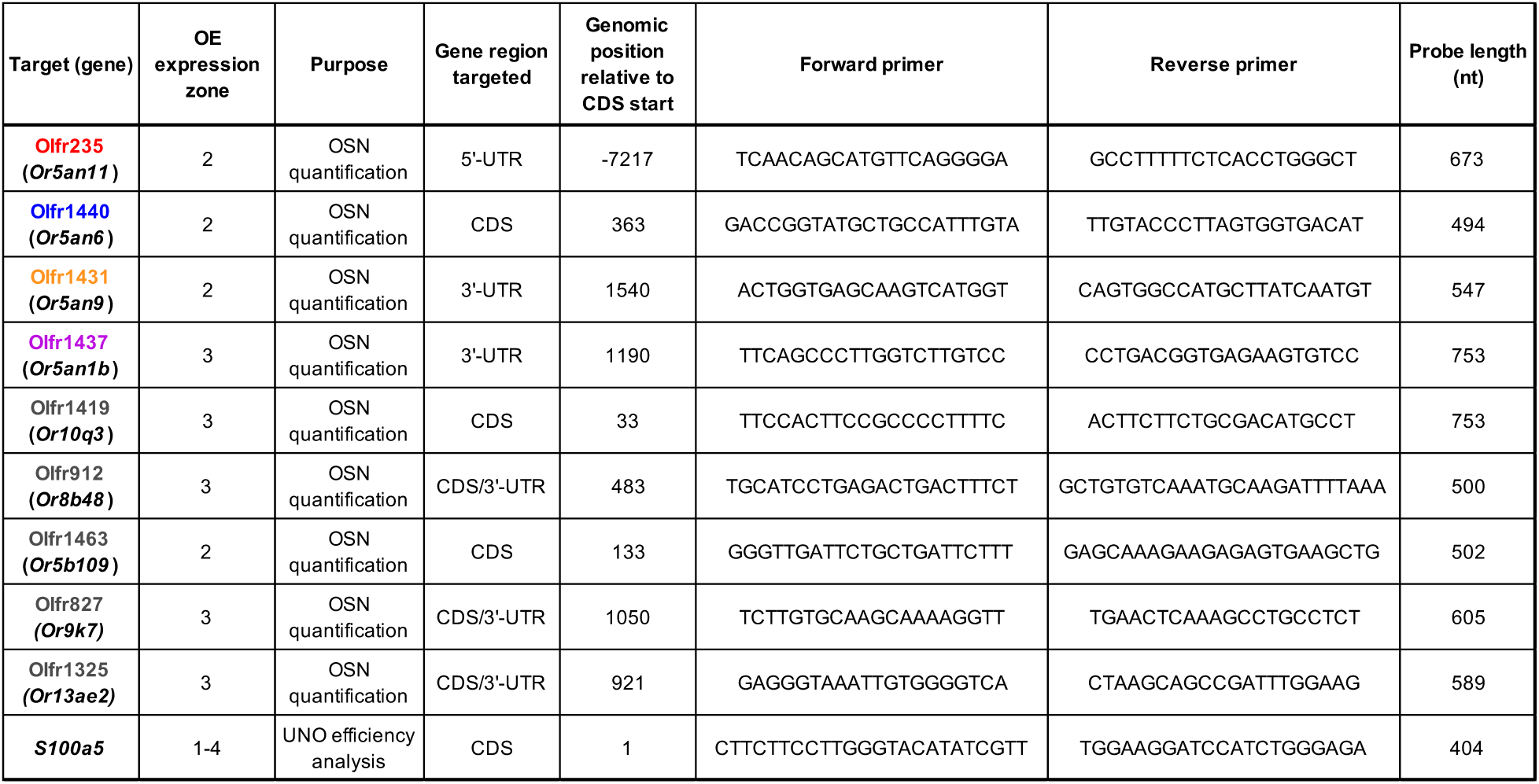
RNA-FISH probes used in this study.

**Appendix 4–table 1.**
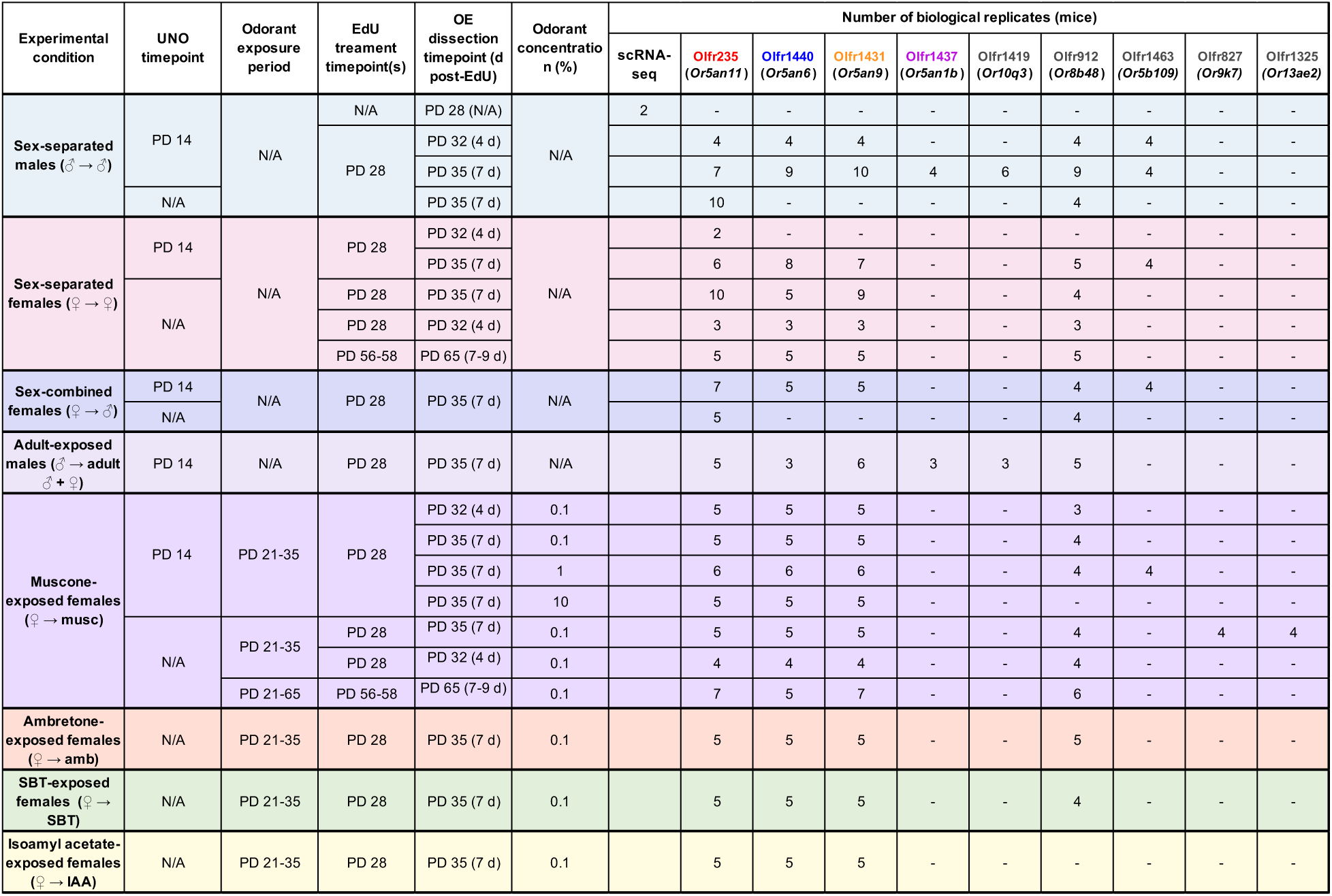
Summary of experimental conditions and numbers of biological replicates (mice) tested for each condition.

**Appendix 4–figure 1.**
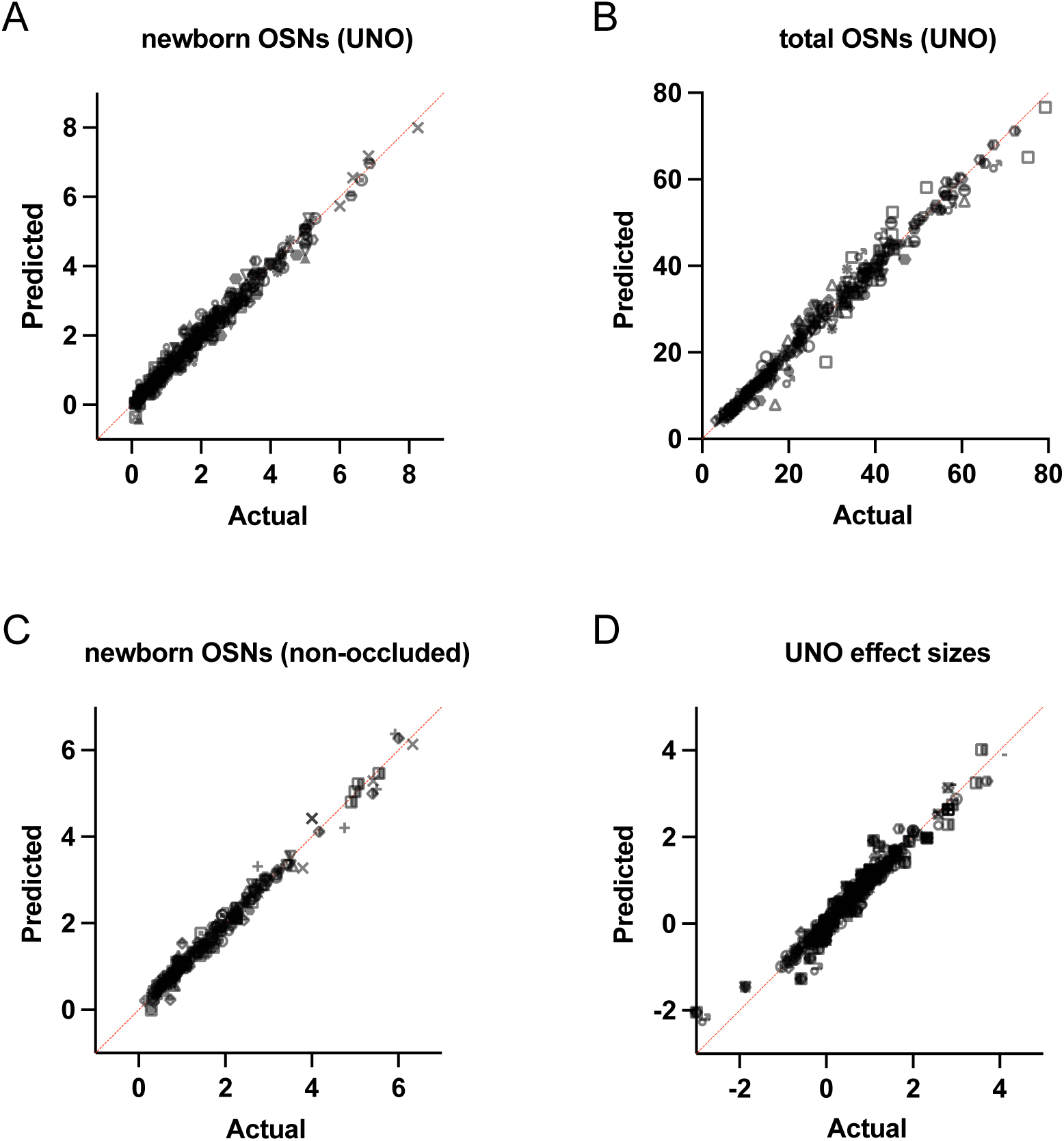
Quantile-quantile (QQ) plots for comparison of actual data to a theoretically normal distribution. A. Analysis of quantities of newborn OSNs on the open and closed sides of the OEs of UNO-treated mice. B. Analysis of quantities of total OSNs on the open and closed sides of the OEs of UNO-treated mice. C. Analysis of quantities of newborn OSNs within the OEs of non-occluded mice. D. Analysis of UNO effect sizes of newborn and total OSNs on the open and closed sides of the OEs of UNO-treated mice. In each plot, predicted values (assuming sampling from a Gaussian distribution) are plotted on the vertical axis, while actual values are plotted on the horizontal axis. Points generally follow the line of identity, indicating that the data reflect a Gaussian (normal) distribution.

## Methods

### Experimental model and subject details

All procedures involving mice were carried out in accordance with NIH standards and approved by the University of Colorado Anschutz Medical Campus Institutional Animal Care and Use Committee (IACUC). For all experiments described, tissue samples were obtained from male or female C57Bl/6J mice that were 28, 35, or 65 days of age at the time of sacrifice. Except for adult-exposed mice, which remained with their parents until sacrifice, all mice were weaned at PD 21 and group-housed either sex-separated (4 females or 4 males per cage) or sex-combined (2 females and 2 males per cage) in standard cages.

### Method details

- <u>Unilateral naris occlusion (UNO)</u>. P14 pups were administered buprenorphine (extended release; 1 mg/kg) subcutaneously using a 31-gauge needle, anesthetized using isoflurane (completeness of anesthesia confirmed through a tail pinch), and then immediately subjected to electrocautery for ∼5 seconds on the right nostril under a dissecting microscope. During electrocautery care was taken to avoid contact of the electrocautery unit with any non-superficial tissues. Pups were examined daily following the procedure to ensure complete blockage of the right nostril through scar formation (typically ∼3–5 days after the procedure) and normal development and activity.
- Preparation of single cells from open and closed sides of OEs of UNO-treated mice for single-cell RNA sequencing (scRNA-seq) analyses. Generation of the OE 2 dataset (this study) was performed as described previously for dataset OE 1^30^, with modifications. Briefly, the OE was dissected from a PD 28 male mouse that had been UNO-treated at PD 14 and weaned sex-separated at PD 21 (Figure 1A, B). The OE was separated at the midline using a clean scalpel and each half was separately minced in 750 µL Hibernate AB Complete media (HAB; BrainBits, HAB100) using a razor blade. For each half, the minced tissue and HAB, along with an additional 750 µL of HAB, were transferred to a 15-mL conical tube using a 1000-μL wide-bore pipette tip and allowed to settle for 1-min, after which the HAB supernatant was removed and transferred to a new 15-mL conical tube. Minced OE halves were each suspended in 1.5 mL of papain solution (2 mg/mL papain [BrainBits; PAP] in Hibernate A-minus Calcium medium [HA-Ca; BrainBits, HACA100]) and incubated for 20 minutes at 37°C, after which the papain solution was removed and discarded. To each digested OE half, the previously-saved HAB supernatant was added, 750-µL at a time, and used to triturate the tissue using a 1000-μl wide-bore pipette tip with 10–15 passes to achieve ∼85% tissue dissociation, after which any large pieces of tissue debris were allowed to settle for ∼20s. Suspended cells were transferred and combined into a new 15-mL conical tube, centrifuged (300 RCF; 3 min, 4°C), resuspended in 1 ml pre-warmed neuronal culturing medium (BrainBits; NbActiv1-100, filtered using a 40-μm Flowmi cell strainer (Bel-Art; H13680-0040), quantified using trypan blue staining and hemocytometry, centrifuged (300 RCF; 3 min, 4°C), resuspended in 1x PBS, stained using the LIVE/DEAD Fixable Aqua Dead Cell Stain Kit (Invitrogen, L34957) according to the manufacturer’s instructions, centrifuged (300 RCF; 3 min, 4°C), resuspended in 1 mL of 1% bovine serum albumin in 1x PBS, centrifuged (300 RCF; 3 min, 4°C), and resuspended in 400 µL of sorting buffer (1x PBS, 1mM EDTA, 25 mM Hepes, pH 7, 1% FBS). Live (non-fluorescent) single cells were sorted using an Astrios Cell Sorter (Beckman Coulter) into a 1.5-mL collection tube containing 20 µL of collection buffer (10% FBS in NbActiv1 media), and used for single cell capture and sequencing.
- <u>Single-cell capture, library preparation, and sequencing</u>. Immediately following sorting, single cells were processed and sequenced within the Genomics Core at the University of Colorado, Anschutz Medical Campus. Following quantification *via* hemocytometry, ∼19,000 cells per sample were targeted for capture using a Chromium X controller (10X Genomics) and used to prepare scRNA-seq libraries using the 3’ HT Kit (10X Genomics) according to the manufacturer’s instructions. Libraries were sequenced to a targeted depth of 40,000 reads/cell using a NovaSeq 6000 instrument (Illumina).
- <u>Odorant exposure</u>. A 1-mL aliquot of 0.1, 1, or 10% odorant solution [(R)-3- methylcyclopentadecanone (muscone; Ambeed, Inc., A275816); 5-Cyclohexadecenone (ambretone; TCI Ltd., C0874; 0.1%); 2-(sec-Butyl)-4,5-dihydrothiazole (SBT; Ambeed, Inc., A578012); or isoamyl acetate (IAA; TCI Ltd., A0033)] in propylene glycol was transferred to a compactly folded piece of absorbent paper (KimTech), which was enclosed within a metal tea ball that was hung within a standard mouse cage. Mice (4 per cage) were housed within odorant-containing cages from weaning (PD 21) until sacrifice (PD 35 or PD 65), with the odorants refreshed every other day.
- <u>2-Deoxy-5-ethynyluridine (EdU) injections</u>. EdU (Carbosynth; NE08701) was administered intraperitoneally to C57Bl/6J mice at PD 28 or PD 56–58 (2 injections/day, spaced 3 h apart, of 10 mg/mL EdU in PBS; 50 mg EdU/kg mouse body weight/injection) ^46^.
- <u>RNA-fluorescent *in situ* hybridization (RNA-FISH) probe design and generation</u>. FISH probes were designed to span 500-1000 base pairs and were targeted to CDS and/or UTR regions of each mRNA transcript (Appendix 3–table 1). Probes were designed to minimize predicted cross-hybridization with off-target transcripts, as assessed using BLAST (NCBI) alignment to the mouse genome. To further minimize off-target cross-hybridization, probes aligning to 3 distinct transcript regions were generated for each target mRNA, and the probe yielding the best signal strength and homogeneity selected for use. Target probe sequences were amplified by PCR from whole OE cDNA using target-specific primers (Appendix 3– table 1), inserted into the pCRII-TOPO vector (ThermoFisher), and confirmed by Sanger sequencing. DIG-labeled antisense riboprobes were generated from 1 µg of linearized plasmid templates using T7 or Sp6 RNA polymerases (NEB) and DIG-11-UTP (Roche), treated with DNaseI (Promega) to remove the template DNA, purified via ethanol precipitation, and dissolved in 30-µL of water. A detailed description of this procedure was published previously ^46^.
- <u>One-color RNA fluorescent *in situ* hybridization (RNA-FISH) combined with EdU staining</u> *<u>via</u>* <u>click chemistry</u>. OEs were dissected from experimental mice (age PD 35 or PD 65), placed in a cryomold containing OCT, flash-frozen in liquid-nitrogen-cooled isopentane, and stored at −80 °C until sectioning. Tissues were cut into 12-μm thick cryo-sections, placed onto slides, and stored at −80 °C until staining. Slide-mounted sections were warmed (37 °C, 5 min), equilibrated in phosphate-buffered saline (PBS; pH 7.2; 3 min, room temperature [RT]), fixed in paraformaldehyde (PFA; 4% in PBS; 10 min, RT), washed in PBS (3 min, RT), permeabilized with Triton-X-100 (0.5% in PBS; 10 min, RT) followed by sodium dodecyl sulfate (1% in PBS; 3 min, RT), washed in PBS (3 × 3 min, RT), incubated in acetylation solution (triethanolamine [0.1 M; pH 7.5], acetic anhydride [0.25%]; 10 min, RT), washed in PBS (3 × 3 min, RT), incubated in hybridization solution (formamide [50%], SSC [5×], Denhardts [5×], yeast tRNA [250 µg/mL], herring sperm DNA [500 µg/mL], heparin [50 µg/mL], EDTA [2.5 mM], Tween-20 [0.1%], CHAPS [0.25 %]; 30 min, RT), hybridized with a DIG-labeled antisense RNA probe (1:750 in hybridization solution; 16 hr, 65 °C), washed with SSC (5×; 1 × 5 min, 65 °C), washed with SSC (0.2×; 4 × 20 min, 65 °C), incubated in H_2_O_2_ (3% in TN [Tris-HCl (0.1 M; pH 7.5), 0.15 M NaCl]; 30 min, RT), washed in TNT (Tween-20 [0.05%] in TN; 5 × 3 min, RT), incubated in TNB (Blocking Reagent [Perkin Elmer; 0.05% in TN]; 30 min, RT), incubated with anti-DIG-POD antibody (Roche; 1:1000 in TNB; 12 hr, 4 °C), and washed in TNT (3 × 20 min, RT). Fluorescent signals corresponding to the target transcript were generated using the Tyramide Signal Amplification (TSA) Plus Fluorescein Kit (Perkin Elmer) according to the manufacturer’s instructions. Slides were washed in 3% BSA in PBS (2 × 5 min, RT, with gentle rocking), incubated with EdU reaction solution (4 mM CuSO_4_, 4 µM Sulfo-Cyanine 3 Azide [Lumiprobe], 100 mM sodium ascorbate [prepared fresh], in PBS; 30 min, RT, in darkness), and washed with 3% BSA in PBS (2 × 3 min, RT). Slides were washed in TNT (2 × 3 min, RT), incubated in DAPI (300 nM in TN; 3 min, RT), washed in TNT (1 × 3 min, RT), and mounted using Vectashield (Vector Laboratories). A detailed description of this procedure was published previously ^46^.
- <u>Image acquisition and processing</u>. Images were acquired using a Zeiss LSM 900 with Airyscan 2 microscope with an automated stage and Zen Blue software (Zeiss). Mosaic images were stitched and each fluorescence channel was adjusted individually to enhance contrast using Zen Blue software. Representative images were exported in jpg format, and rotated and cropped using Adobe Photoshop. A detailed description of this procedure was published previously ^46^.
- <u>Quality criteria for sectioned OEs</u>. For UNO-treated mice, UNO efficiency was assessed by visual inspection of mouse nostrils prior to dissection. Additionally, OE cryosections from UNO-treated mice were stained *via* one-color RNA-FISH for *S100a5* transcript levels. For each section analyzed, *S100a5* mRNA intensities were evaluated within paired regions on the two sides of each OE section. OEs from UNO-treated mice were excluded from further analysis if the mean fluorescence intensity corresponding to *S100a5* transcripts was not noticeably (∼10-fold) greater on the open side of the OE compared to the closed side. All sections from both UNO-treated and non-occluded mice were assessed for left-right symmetry and intactness. Individual OE sections were excluded if they lacked a high degree of symmetry or were less than 90% intact. No sections were otherwise excluded. A detailed description of this procedure was published previously ^46^.
- <u>Gas chromatography – mass spectrometry (GC-MS) analysis of mouse preputial gland</u> <u>(PG) extracts</u>. Immediately following euthanasia, three PGs were dissected from 5-week-old male mice, transferred to separate 1.5-mL microcentrifuge tubes, and flash-frozen on dry ice. Frozen PG tissues were shipped on dry ice to the Analytical Resource Core at Colorado State University for processing, essentially as described previously ^79^. Each gland was ground in liquid nitrogen using a mortar and pestle, transferred to a glass extraction vial containing 200 µL of dichloromethane, vortexed and sonicated for 30 min, and centrifugated at 3,000 g for 1 min. Supernatants were transferred to glass autosampler vials and stored at −20 °C until analysis. For GC-MS analyses, 1 μL of dichloromethane extract was injected into a Trace 1310 gas chromatography instrument coupled to a Thermo ISQ-LT mass spectrometer, at a 5:1 split ratio. The inlet was held at 285 °C. Separation was achieved on a 30m DB-5MS column (J&W, 0.25 mm ID, 0.25 µm film thickness). Oven temperature was held at 80 °C for 0.5 min, ramped at 15°C/min to 330°C, and held at 330 °C for 8 min. Helium carrier gas flow was held at 1.2 mL/min. Temperatures of transfer line and ion source were held at 300 °C and 260 °C, respectively. Fullscan mode (50-650 m/z) was used on all the samples. The injection order of samples was randomized. Standards: (R)-3-Methylcyclopentadecanone (Muscone) was purchased from Ambeed, Inc. (A275816), 5-Cyclohexadecenone (Ambretone) was from TCI America (C0874), 8-Cyclohexadecenone (Globanone) was from Perfumer’s Apprentice (ac-1510-sz1), and Cyclopentadecanol was from TCI America (C1528). GC-MS data were processed using Chromeleon 7.2.10 software (Thermo Scientific).

### Quantification and statistical analysis

- <u>scRNA-seq analysis of UNO-induced changes in subtype-specific quantities of newborn</u> <u>OSNs</u>. Newborn OSNs of specific subtypes were quantified within scRNA-seq datasets generated from the open and closed sides of the OEs of 2 different male C57Bl/6J mice that were UNO-treated at PD 14 and sacrificed at PD 28 (Figure 1): OE 1 (generated previously ^30^; https://www.ncbi.nlm.nih.gov/geo/query/acc.cgi?acc=GSE157119), and OE 2 (this study; https://www.ncbi.nlm.nih.gov/geo/query/acc.cgi?acc=GSE278693). Newborn OSNs of specific subtypes were identified by their expression of the iOSN-specific marker *Gap43* (Log_2_ UMI > 1) and a specific OR gene (Log_2_ UMI > 2) (Figure 1) and quantified within the open and closed sides of the OE 1 and OE 2 datasets using Loupe Browser software (8.0.0; 10X Genomics). Subtype-specific quantities of newborn OSNs in each OE half were normalized based on the total number of cells or OSNs (*Gap43* Log_2_ UMI > 1 or *Omp* Log_2_ UMI > 3) within the corresponding dataset.
- scRNA-seq analysis of OR transcripts within individual INP3, iOSN, and mOSN cells on the open and closed sides of the OEs of UNO-treated mice. Individual INP3, iOSN, and mOSN cells expressing transcripts of a specific OR gene were identified based on their expression of *Tex15* ^38,43^ (Log_2_ UMI > 2) and the OR (Log_2_ UMI > 0), their expression of *Gap43* (Log_2_ UMI > 1) and the OR (Log_2_ UMI > 2), or their expression the OR (Log_2_ UMI > 2) and lack of expression of *Gap43* (Log_2_ UMI ≤ 1), respectively, using Loupe Browser software (8.0.0). The cellular transcript levels of all ORs within INP3, iOSN, and mOSN cells expressing a specific musk-responsive (Olfr235, Olfr1440) or non-musk responsive (Olfr701) OR within the open and closed datasets of OE 1 and OE 2 samples were calculated and exported using Loupe Browser software (8.0.0).
- <u>Histological quantification of newborn and total OSNs of specific subtypes</u>. Quantities of newborn and total OSNs within the OEs of individual mice were determined from images of a series of at least 5 coronal tissue sections stained for EdU and a specific OR mRNA and spanning the anterior-posterior length of each OE. Quantifications were performed separately on the two sides of each OE section, with the experimenter blinded to the sample identities. Newborn cells of specific subtypes were identified based on robust nuclear EdU staining (Cy3) (> 2-fold above background) that was overlapped at least 50% by OR mRNA staining (FITC). A detailed description of this procedure was published previously ^46^.
- <u>Comparison of methods for normalizing quantities of newborn OSNs of specific subtypes</u>. For all histological quantifications of newborn and total OSNs of specific subtypes in this study, OSN quantities were normalized to the number of half-sections analyzed. This approach has been used in multiple previous studies for quantifying newborn (OR+/EdU+) and total (OR+) OSN abundances ^17,19,30,46^. To verify the rigor of this method, newborn Olfr235 OSN quantities were also normalized by total quantities of newborn OSNs (approximated by total EdU+ cells) or total quantities of cells (approximated by DAPI+ area) for three experimental comparisons in this study: the open versus the closed sides of OEs from UNO-treated male mice (Figure 2–figure supplement 2), the OEs from non-occluded females versus males (Figure 3–figure supplement 2), and the OEs of non-occluded females exposed just to female littermates versus females also exposed to 0.1% muscone (Figure 4– figure supplement 4). For each condition, EdU+ cells and DAPI+ areas were identified and quantified within images of OE sections stained for Olfr235 mRNA and EdU using the surface and stats tools, respectively, within Imaris software (version 10.2).
- <u>Statistics</u>. To assess deviation from normality of the histological quantifications of newborn and total OSNs of specific subtypes used for comparisons in this study, all datasets were tested using the Shapiro-Wilk test for non-normality and the P values obtained are included within Supplementary file 1. Of the 274 datasets tested, 253 have Shapiro-Wilk P values > 0.05, indicating that the vast majority (92%) do not show evidence of significant deviation from a normal distribution. A general lack of deviation of the datasets in this study from a normal distribution is further supported by quantile-quantile (QQ) plots (Appendix 4–figure 1). Moreover, results of both parametric and non-parametric statistical tests of comparisons in this study are, in general, in good agreement (Supplementary file 2). Statistical analyses of differences in OSN quantities between the open and closed sides of OEs from UNO-treated mice were performed using ratio paired two-tailed *t*-tests (parametric) and Wilcoxon matched-pairs signed rank two-tailed tests (non-parametric). For these tests, the open and closed sides of each OE were paired, enabling testing of differences between the two sides independent of OSN number and staining variance between sections. Statistical analyses of differences in OSN quantities in the OEs of non-occluded mice or UNO effect sizes in UNO- treated mice subjected to two different experimental conditions were performed using unpaired two-tailed *t*-tests (parametric) and Mann-Whitney two-tailed tests (non-parametric). Statistical analyses of differences in OSN quantities in the OEs of non-occluded mice or UNO effect sizes in UNO-treated mice subjected more than two different experimental conditions were performed using one-way ANOVA tests, FDR-adjusted using the 2-stage linear step-up procedure of Benjamini, Krieger and Yekutieli (parametric) and Kruskal-Wallis tests, FDR-adjusted using the 2-stage linear step-up procedure of Benjamini, Krieger and Yekutieli (non-parametric). For comparisons of differences in quantities of newborn OSNs of musk-responsive subtypes at 4 and 7 days post-EdU between non-occluded mice exposed and unexposed to muscone, a two sample ANOVA - fixed-test, using F distribution (right-tailed) was used. Data presented in figures represent mean +/- SEM. For all statistical analyses, a significance threshold of *P* < 0.05 was used. All statistical tests were performed using Prism 10 software (Graphpad). Results of all statistical tests performed in this study are summarized in Supplementary file 2. P values reported within the text and figures are based on parametric tests.
- <u>Sample-size estimation</u>. Results from previous studies ^19,30^ were used to determine an appropriate sample size for comparing the number of total (OR+) and newborn (OR+/EdU+) OSNs on the open and closed sides of the OE. Previously, it was found that for an OR with a typical expression frequency (∼0.1%) and an effect size of ∼2-fold, 12 OE sections taken from four different animals were sufficient to find a highly statistically significant difference (*P* < 0.001; paired two-tailed *t* test). For comparisons between different animals, results from previous studies ^19,30^ were also used to determine an appropriate sample size. Previously, we had found that for an OR with a typical expression frequency (∼0.1%) and an effect size of ∼2-fold, 20 OE sections taken from four different animals was sufficient to find a highly statistically significant difference between different animals (*P* < 0.01; two-tailed unpaired *t* test).

## References

1 Imai T. Development of the Olfactory System. Evol. Neurosensory Cells Syst. 1st ed. Boca Raton: CRC Press; 2022. p. 29–60.

2 Xu L, Li W, Voleti V, Zou D-J, Hillman EMC, Firestein S. Widespread receptor-driven modulation in peripheral olfactory coding. Science 2020;368:eaaz5390. 10.1126/science.aaz5390.

3 Brann JH, Firestein SJ. A lifetime of neurogenesis in the olfactory system. Front Neurosci 2014;8:182. 10.3389/fnins.2014.00182.

4 Schwob JE, Jang W, Holbrook EH, Lin B, Herrick DB, Peterson JN, et al. Stem and progenitor cells of the mammalian olfactory epithelium: Taking poietic license. J Comp Neurol 2017;525:1034–54. 10.1002/cne.24105.

5 Yu CR, Wu Y. Regeneration and rewiring of rodent olfactory sensory neurons. Exp Neurol 2017;287:395–408. 10.1016/j.expneurol.2016.06.001.

6 Lledo P-M, Valley M. Adult Olfactory Bulb Neurogenesis. Cold Spring Harb Perspect Biol 2016;8:. 10.1101/cshperspect.a018945.

7 Ming G-L, Song H. Adult neurogenesis in the mammalian brain: significant answers and significant questions. Neuron 2011;70:687–702. 10.1016/j.neuron.2011.05.001.

8 Opendak M, Gould E. Adult neurogenesis: a substrate for experience-dependent change. Trends Cogn Sci 2015;19:151–61. 10.1016/j.tics.2015.01.001.

9 McClintock TS. Odorant Receptor Gene Choice. ChemoSense 2015;16:3–13.

10 Monahan K, Lomvardas S. Monoallelic expression of olfactory receptors. Annu Rev Cell Dev Biol 2015;31:721–40. 10.1146/annurev-cellbio-100814-125308.

11 Yusuf N, Monahan K. Epigenetic programming of stochastic olfactory receptor choice. Genes N Y N 2000 2024;62:e23593. 10.1002/dvg.23593.

12 Cadiou H, Aoudé I, Tazir B, Molinas A, Fenech C, Meunier N, et al. Postnatal odorant exposure induces peripheral olfactory plasticity at the cellular level. J Neurosci Off J Soc Neurosci 2014;34:4857–70. 10.1523/JNEUROSCI.0688-13.2014.

13 Cavallin MA, Powell K, Biju KC, Fadool DA. State-dependent sculpting of olfactory sensory neurons is attributed to sensory enrichment, odor deprivation, and aging. Neurosci Lett 2010;483:90–5. 10.1016/j.neulet.2010.07.059.

14 Coppola DM, Waggener CT. The effects of unilateral naris occlusion on gene expression profiles in mouse olfactory mucosa. J Mol Neurosci MN 2012;47:604–18. 10.1007/s12031-011-9690-4.

15 Dias BG, Ressler KJ. Parental olfactory experience influences behavior and neural structure in subsequent generations. Nat Neurosci 2014;17:89–96. 10.1038/nn.3594.

16 Fischl AM, Heron PM, Stromberg AJ, McClintock TS. Activity-dependent genes in mouse olfactory sensory neurons. Chem Senses 2014;39:439–49. 10.1093/chemse/bju015.

17 Ibarra-Soria X, Nakahara TS, Lilue J, Jiang Y, Trimmer C, Souza MA, et al. Variation in olfactory neuron repertoires is genetically controlled and environmentally modulated. eLife 2017;6:. 10.7554/eLife.21476.

18 Jones SV, Choi DC, Davis M, Ressler KJ. Learning-dependent structural plasticity in the adult olfactory pathway. J Neurosci Off J Soc Neurosci 2008;28:13106–11. 10.1523/JNEUROSCI.4465-08.2008.

19 van der Linden C, Jakob S, Gupta P, Dulac C, Santoro SW. Sex separation induces differences in the olfactory sensory receptor repertoires of male and female mice. Nat Commun 2018;9:5081. 10.1038/s41467-018-07120-1.

20 Morrison FG, Dias BG, Ressler KJ. Extinction reverses olfactory fear-conditioned increases in neuron number and glomerular size. Proc Natl Acad Sci U S A 2015;112:12846–51. 10.1073/pnas.1505068112.

21 Santoro SW, Dulac C. The activity-dependent histone variant H2BE modulates the life span of olfactory neurons. eLife 2012;1:e00070. 10.7554/eLife.00070.

22 Watt WC, Sakano H, Lee ZY, Reusch JE, Trinh K, Storm DR. Odorant stimulation enhances survival of olfactory sensory neurons via MAPK and CREB. Neuron 2004;41:955–67. S0896627304000753 [pii].

23 Zhao S, Tian H, Ma L, Yuan Y, Yu CR, Ma M. Activity-Dependent Modulation of Odorant Receptor Gene Expression in the Mouse Olfactory Epithelium. PLoS ONE 2013;8:e69862. 10.1371/journal.pone.0069862.

24 Vihani A, Hu XS, Gundala S, Koyama S, Block E, Matsunami H. Semiochemical responsive olfactory sensory neurons are sexually dimorphic and plastic. eLife 2020;9:e54501. 10.7554/eLife.54501.

25 Ross JM, Fletcher ML. Aversive learning-induced plasticity throughout the adult mammalian olfactory system: insights across development. J Bioenerg Biomembr 2018. 10.1007/s10863-018-9770-z.

26 Zhao H, Reed RR. X inactivation of the OCNC1 channel gene reveals a role for activity-dependent competition in the olfactory system. Cell 2001;104:651–60. 10.1016/s0092-8674(01)00262-8.

27 Farbman AI, Brunjes PC, Rentfro L, Michas J, Ritz S. The effect of unilateral naris occlusion on cell dynamics in the developing rat olfactory epithelium. J Neurosci Off J Soc Neurosci 1988;8:3290–5.

28 Cummings DM, Brunjes PC. Changes in cell proliferation in the developing olfactory epithelium following neonatal unilateral naris occlusion. Exp Neurol 1994;128:124–8. 10.1006/exnr.1994.1119.

29 Mirich JM, Brunjes PC. Activity modulates neuronal proliferation in the developing olfactory epithelium. Brain Res Dev Brain Res 2001;127:77–80.

30 van der Linden CJ, Gupta P, Bhuiya AI, Riddick KR, Hossain K, Santoro SW. Olfactory Stimulation Regulates the Birth of Neurons That Express Specific Odorant Receptors. Cell Rep 2020;33:108210. 10.1016/j.celrep.2020.108210.

31 Coppola DM. Studies of olfactory system neural plasticity: the contribution of the unilateral naris occlusion technique. Neural Plast 2012;2012:351752. 10.1155/2012/351752.

32 McClintock TS, Adipietro K, Titlow WB, Breheny P, Walz A, Mombaerts P, et al. In vivo identification of eugenol-responsive and muscone-responsive mouse odorant receptors. J Neurosci Off J Soc Neurosci 2014;34:15669–78. 10.1523/JNEUROSCI.3625-14.2014.

33 Sato-Akuhara N, Horio N, Kato-Namba A, Yoshikawa K, Niimura Y, Ihara S, et al. Ligand Specificity and Evolution of Mammalian Musk Odor Receptors: Effect of Single Receptor Deletion on Odor Detection. J Neurosci Off J Soc Neurosci 2016;36:4482–91. 10.1523/JNEUROSCI.3259-15.2016.

34 Shirasu M, Yoshikawa K, Takai Y, Nakashima A, Takeuchi H, Sakano H, et al. Olfactory receptor and neural pathway responsible for highly selective sensing of musk odors. Neuron 2014;81:165–78. 10.1016/j.neuron.2013.10.021.

35 Peterlin Z, Firestein S, Rogers ME. The state of the art of odorant receptor deorphanization: a report from the orphanage. J Gen Physiol 2014;143:527–42. 10.1085/jgp.201311151.

36 McIntyre JC, Titlow WB, McClintock TS. Axon growth and guidance genes identify nascent, immature, and mature olfactory sensory neurons. J Neurosci Res 2010;88:3243–56. 10.1002/jnr.22497.

37 Verhaagen J, Oestreicher AB, Gispen WH, Margolis FL. The expression of the growth associated protein B50/GAP43 in the olfactory system of neonatal and adult rats. J Neurosci Off J Soc Neurosci 1989;9:683–91.

38 Fletcher RB, Das D, Gadye L, Street KN, Baudhuin A, Wagner A, et al. Deconstructing Olfactory Stem Cell Trajectories at Single-Cell Resolution. Cell Stem Cell 2017;20:817–830.e8. 10.1016/j.stem.2017.04.003.

39 Tan L, Xie XS. A Near-Complete Spatial Map of Olfactory Receptors in the Mouse Main Olfactory Epithelium. Chem Senses 2018;43:427–32. 10.1093/chemse/bjy030.

40 Hanchate NK, Kondoh K, Lu Z, Kuang D, Ye X, Qiu X, et al. Single-cell transcriptomics reveals receptor transformations during olfactory neurogenesis. Science 2015;350:1251–5. 10.1126/science.aad2456.

41 Tan L, Li Q, Xie XS. Olfactory sensory neurons transiently express multiple olfactory receptors during development. Mol Syst Biol 2015;11:844. 10.15252/msb.20156639.

42 Bashkirova EV, Klimpert N, Monahan K, Campbell CE, Osinski J, Tan L, et al. Opposing, spatially-determined epigenetic forces impose restrictions on stochastic olfactory receptor choice. eLife 2023;12:RP87445. 10.7554/eLife.87445.

43 Pourmorady AD, Bashkirova EV, Chiariello AM, Belagzhal H, Kodra A, Duffié R, et al. RNA-mediated symmetry breaking enables singular olfactory receptor choice. Nature 2024;625:181–8. 10.1038/s41586-023-06845-4.

44 Saraiva LR, Ibarra-Soria X, Khan M, Omura M, Scialdone A, Mombaerts P, et al. Hierarchical deconstruction of mouse olfactory sensory neurons: from whole mucosa to single-cell RNA-seq. Sci Rep 2015;5:18178. 10.1038/srep18178.

45 Scholz P, Kalbe B, Jansen F, Altmueller J, Becker C, Mohrhardt J, et al. Transcriptome Analysis of Murine Olfactory Sensory Neurons during Development Using Single Cell RNA-Seq. Chem Senses 2016;41:313–23. 10.1093/chemse/bjw003.

46 Hossain K, Smith M, Santoro SW. A histological protocol for quantifying the birthrates of specific subtypes of olfactory sensory neurons in mice. STAR Protoc 2023;4:102432. 10.1016/j.xpro.2023.102432.

47 Osada K, Tashiro T, Mori K, Izumi H. The identification of attractive volatiles in aged male mouse urine. Chem Senses 2008;33:815–23. 10.1093/chemse/bjn045.

48 Osada K, Yamazaki K, Curran M, Bard J, Smith BPC, Beauchamp GK. The scent of age. Proc Biol Sci 2003;270:929–33. 10.1098/rspb.2002.2308.

49 Schwende FJ, Wiesler D, Jorgenson JW, Carmack M, Novotny M. Urinary volatile constituents of the house mouse,Mus musculus, and their endocrine dependency. J Chem Ecol 1986;12:277–96. 10.1007/BF01045611.

50 Stopková R, Matějková T, Dodoková A, Talacko P, Zacek P, Sedlacek R, et al. Variation in mouse chemical signals is genetically controlled and environmentally modulated. Sci Rep 2023;13:8573. 10.1038/s41598-023-35450-8.

51 Kelemen G. The junction of the nasal cavity and the pharyngeal tube in the rat. Arch Otolaryngol 1947;45:159–68. 10.1001/archotol.1947.00690010168002.

52 Shykind BM, Rohani SC, O’Donnell S, Nemes A, Mendelsohn M, Sun Y, et al. Gene switching and the stability of odorant receptor gene choice. Cell 2004;117:801–15. 10.1016/j.cell.2004.05.015.

53 Rodriguez-Gil DJ, Bartel DL, Jaspers AW, Mobley AS, Imamura F, Greer CA. Odorant receptors regulate the final glomerular coalescence of olfactory sensory neuron axons. Proc Natl Acad Sci U S A 2015;112:5821–6. 10.1073/pnas.1417955112.

54 Han X, Jiang Y, Feng N, Yang P, Zhang M, Jin W, et al. Comparison of the Homology Between Muskrat Scented Gland and Mouse Preputial Gland. J Mamm Evol 2022;29:435–46. 10.1007/s10914-022-09604-w.

55 Mookherjee BD, Wilson RA. The chemistry and fragrance of natural musk compounds. T Theimer E Ed Fragr Chem Sci Sense Smell 2012:433.

56 Suh KS, Kim SY, Bae YC, Ronnett GV, Moon C. Effects of unilateral naris occlusion on the olfactory epithelium of adult mice. Neuroreport 2006;17:1139–42. 10.1097/01.wnr.0000224762.54336.7d.

57 van Drongelen W, Holley A, Døving KB. Convergence in the olfactory system: quantitative aspects of odour sensitivity. J Theor Biol 1978;71:39–48. 10.1016/0022-5193(78)90212-6.

58 Meisami E. A proposed relationship between increases in the number of olfactory receptor neurons, convergence ratio and sensitivity in the developing rat. Brain Res Dev Brain Res 1989;46:9–19. 10.1016/0165-3806(89)90139-9.

59 D’Hulst C, Mina RB, Gershon Z, Jamet S, Cerullo A, Tomoiaga D, et al. MouSensor: A Versatile Genetic Platform to Create Super Sniffer Mice for Studying Human Odor Coding. Cell Rep 2016;16:1115–25. 10.1016/j.celrep.2016.06.047.

60 Apfelbach R, Russ D, Slotnick BM. Ontogenetic changes in odor sensitivity, olfactory receptor area and olfactory receptor density in the rat. Chem Senses 1991;16:209–18. 10.1093/chemse/16.3.209.

61 Wang HW, Wysocki CJ, Gold GH. Induction of olfactory receptor sensitivity in mice. Science 1993;260:998–1000. 10.1126/science.8493539.

62 Yee KK, Wysocki CJ. Odorant exposure increases olfactory sensitivity: olfactory epithelium is implicated. Physiol Behav 2001;72:705–11. 10.1016/s0031-9384(01)00428-0.

63 Dalton P, Doolittle N, Breslin PAS. Gender-specific induction of enhanced sensitivity to odors. Nat Neurosci 2002;5:199–200. 10.1038/nn803.

64 Voznessenskaya V, Parfyonova V, Wysocki C. Induced olfactory sensitivity in rodents: a general phenomenon. Adv Biosci 1995;93:399–406.

65 Wang L, Chen L, Jacob T. Evidence for peripheral plasticity in human odour response. J Physiol 2004;554:236–44. 10.1113/jphysiol.2003.054726.

66 Wysocki CJ, Dorries KM, Beauchamp GK. Ability to perceive androstenone can be acquired by ostensibly anosmic people. Proc Natl Acad Sci U S A 1989;86:7976–8. 10.1073/pnas.86.20.7976.

67 Zou D-J, Chesler A, Firestein S. How the olfactory bulb got its glomeruli: a just so story? Nat Rev Neurosci 2009;10:611–8. 10.1038/nrn2666.

68 Yamada Y, Bhaukaurally K, Madarász TJ, Pouget A, Rodriguez I, Carleton A. Context- and Output Layer-Dependent Long-Term Ensemble Plasticity in a Sensory Circuit. Neuron 2017;93:1198–1212.e5. 10.1016/j.neuron.2017.02.006.

69 Dorrego-Rivas A, Grubb MS. Developing and maintaining a nose-to-brain map of odorant identity. Open Biol 2022;12:220053. 10.1098/rsob.220053.

70 Qiu Q, Wu Y, Ma L, Xu W, Hills M, Ramalingam V, et al. Acquisition of innate odor preference depends on spontaneous and experiential activities during critical period. eLife 2021;10:e60546. 10.7554/eLife.60546.

71 Asada R, Kageyama K, Tanaka H, Saitoh Y, Miwa N. Antitumor and anti-invasive effects of diverse musk-fragrant macrocyclic ketones and their enhancement by hyperthermia. Mol Med Rep 2012;5:148–52.

72 Ruzicka L, Stoll M, Schinz H. Zur Kenntnis des Kohlenstoffringes II. Synthese der carbocyclischen Ketone vom Zehner-bis zum Achtzehnerring. Helv Chim Acta 1926;9:249– 64.

73 Ward JP, van Dorp DA. The animal musks and a comment of their biogenesis. Experientia 1981;37:917–22.

74 Agosta WC. Chemical communication: the language of pheromones. Henry Holt and Company; 1992.

75 Fukui H, Komaki R, Okui M, Toyoshima K, Kuda K. The effects of odor on cortisol and testosterone in healthy adults. Neuroendocrinol Lett 2007;28:433–7.

76 Kato M, Yamaoka Y, Tanaka H. Study on the stimulation on the secretion of the female sex hormone by some perfumery raw materials. AROMA Res 2004;5:64–8.

77 Horio N, Murata K, Yoshikawa K, Yoshihara Y, Touhara K. Contribution of individual olfactory receptors to odor-induced attractive or aversive behavior in mice. Nat Commun 2019;10:209. 10.1038/s41467-018-07940-1.

78 Santoro SW, Jakob S. Gene expression profiling of the olfactory tissues of sex-separated and sex-combined female and male mice. Sci Data 2018;5:180260. 10.1038/sdata.2018.260.

79 Bala B, Babbar BK. Sex pheromones of Preputial gland: Their chemical identification and role in social &amp; reproductive behaviour of house rat, Rattus rattus 2023. 10.21203/rs.3.rs-2745121/v1.

